# Enterococcal linear plasmids adapt to *Enterococcus faecium* and spread within multidrug-resistant clades

**DOI:** 10.1101/2022.09.07.506885

**Authors:** Yusuke Hashimoto, Masato Suzuki, Sae Kobayashi, Yuki Hirahara, Takahiro Nomura, Jun Kurushima, Hidetada Hirakawa, Koichi Tanimoto, Haruyoshi Tomita

## Abstract

Antimicrobial resistance (AMR) of bacterial pathogens, including enterococci, is a global concern, and plasmids are crucial for spreading and maintaining AMR genes. Plasmids with linear topology were recently identified in clinical multidrug-resistant enterococci. The enterococcal linear-form plasmids, such as pELF1, confer resistance to clinically important antimicrobials, including vancomycin; however, little information exists about their epidemiological and physiological effects. In this study, we identified several lineages of enterococcal linear plasmids that are structurally conserved and occur globally. pELF1-like linear plasmids show plasticity in acquiring and maintaining AMR genes, often via transposition with the mobile genetic element IS*1216E*. This linear plasmid family has several characteristics enabling long-term persistence in the bacterial population, including high horizontal self-transmissibility, low-level transcription of plasmid-encoded genes, and a moderate effect on the *Enterococcus faecium* genome alleviating fitness cost and promoting vertical inheritance. Combined with its broad host range, the linear plasmid is an important factor in the spread and maintenance of AMR genes among enterococci.

## Introduction

Enterococcus faecium, one of “the ESKAPE (<u>E</u>nterococcus faecium, <u>S</u>taphylococcus aureus, <u>K</u>lebsiella pneumoniae, <u>A</u>cinetobacter baumannii, <u>P</u>seudomonas aeruginosa, and <u>E</u>nterobacter species) bugs,” is a major cause of nosocomial infection and frequently presents with multidrug resistance^1^. Vancomycin resistance in enterococci is a serious clinical problem owing to the paucity of treatment options. Vancomycin-resistant E. faecium (VREfm) is the major species among vancomycin-resistant enterococci (VRE)^2^. The propagation of VREfm involves both the spread of clones (i.e., clonal complex 17 [CC17]) and mobile genetic elements (MGEs), such as plasmids containing vancomycin-resistant genes^3, 4^. Particularly for E. faecium, conjugative plasmids are the most important reservoirs and vehicles for vancomycin resistance genes^5, 6^. A comprehensive understanding of enterococcal plasmids, which confer various ecological characteristics, is therefore of great clinical value.

In 2019, we reported an enterococcal linear-form plasmid (pELF) harboring vancomycin resistance gene clusters from a Japanese VRE isolate^7^. In addition to the nosocomial transmission of VRE through a pELF1-like plasmid, pELF2, reports have confirmed the importance of the linear plasmid in the regional spread of VRE^8, 9^. The host range is assumed to be broad, and the self-transmissibility of pELF to enterococci has been confirmed experimentally and clinically^7, 8^. The efficient propagation of drug resistance genes complicates both the treatment and control of disease transmission; thus, to improve clinical outcomes, it is essential that we elucidate the dynamics of plasmid spread. However, the reports and epidemiological data of pELF1-like plasmids are scant, with no information on their biological effects on host enterococci. Here we reported the detection of several pELF1-like plasmid-carrying strains from clinical isolates in Japan. Along with pELF1-like plasmid resources in a public database, we performed an in-depth molecular epidemiology analysis based on whole-genome sequencing (WGS). Furthermore, we analyzed the impact of pELF1-like plasmids on host enterococci fitness at the transcriptional level. This study provides an integrated characterization of enterococcal linear plasmids at the epidemiologic, phenomic, and transcriptomic levels.

## Results

### Prevalence of pELF1-like plasmids in clinical enterococcal isolates in Japan

We analyzed a total of 1,769 strains of VRE (385 strains) and vancomycin-susceptible enterococci (VSE, 1,384 strains, Table 1). VRE strains were isolated in Japan since the early 2000s, whereas VSE strains were isolated from October 2009 to May 2018 at a single core medical institute in Gunma prefecture.

**Table 1.**
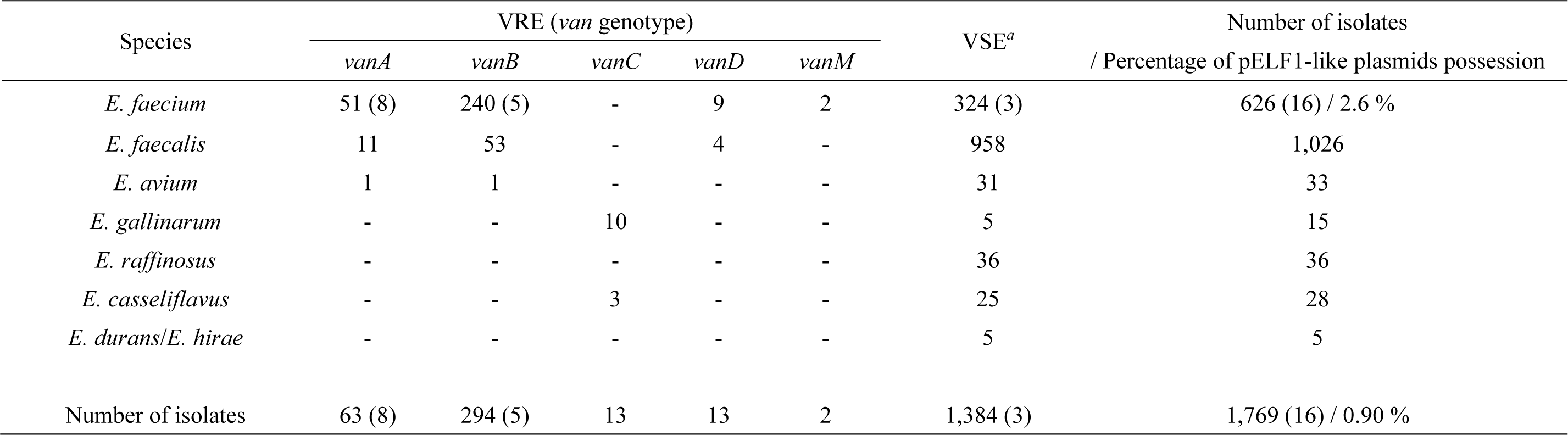
Enterococcal strains for detection of pELF1-like plasmids. Parentheses indicate the number of strains carrying pELF1-like linear plasmids. *^a^*Strains that do not show vancomycin resistance as a phenotype.

Identification of the plasmids was performed via colony PCR using primers designed for paired-end detection of pELF1-like plasmids (Supplementary Table 1). Through PCR screening, pELF1-like plasmids were identified in 16 strains (0.9 % of the total strains), all of which were *E. faecium* (2.6 %, Table 1); 3 of the 1,384 VSE strains (0.2 %) and 13 of the 385 VRE strains (3.4 %) harbored pELF1-like plasmids. Of the thirteen VRE strains, eight were of the VanA-type and five of the VanB-type.

### Characteristics of the pELF1-like plasmid-carrying strains

We characterized the 18 pELF1-like plasmid-carrying strains, including AA708 (pELF1) and KUHS13 (pELF2, Table 2), based on the available epidemiological information. VRE strains carrying the pELF1-like plasmid were detected nationwide (Supplementary Fig. 1A). Among the thirteen pELF1-like plasmids carrying VRE strains, six strains (JHP9, JHP10, JHP35, JHP36, JHP38, and JHP80) had been isolated in the early 2000s. The pELF1-like plasmid was also detected in the seven VRE strains isolated after 2010. Three pELF1-like plasmid-carrying VSE strains were detected in 2014 and 2015, and two of these strains (GK923 and GK961) were detected on the same patient on different days.

**Table 2.**
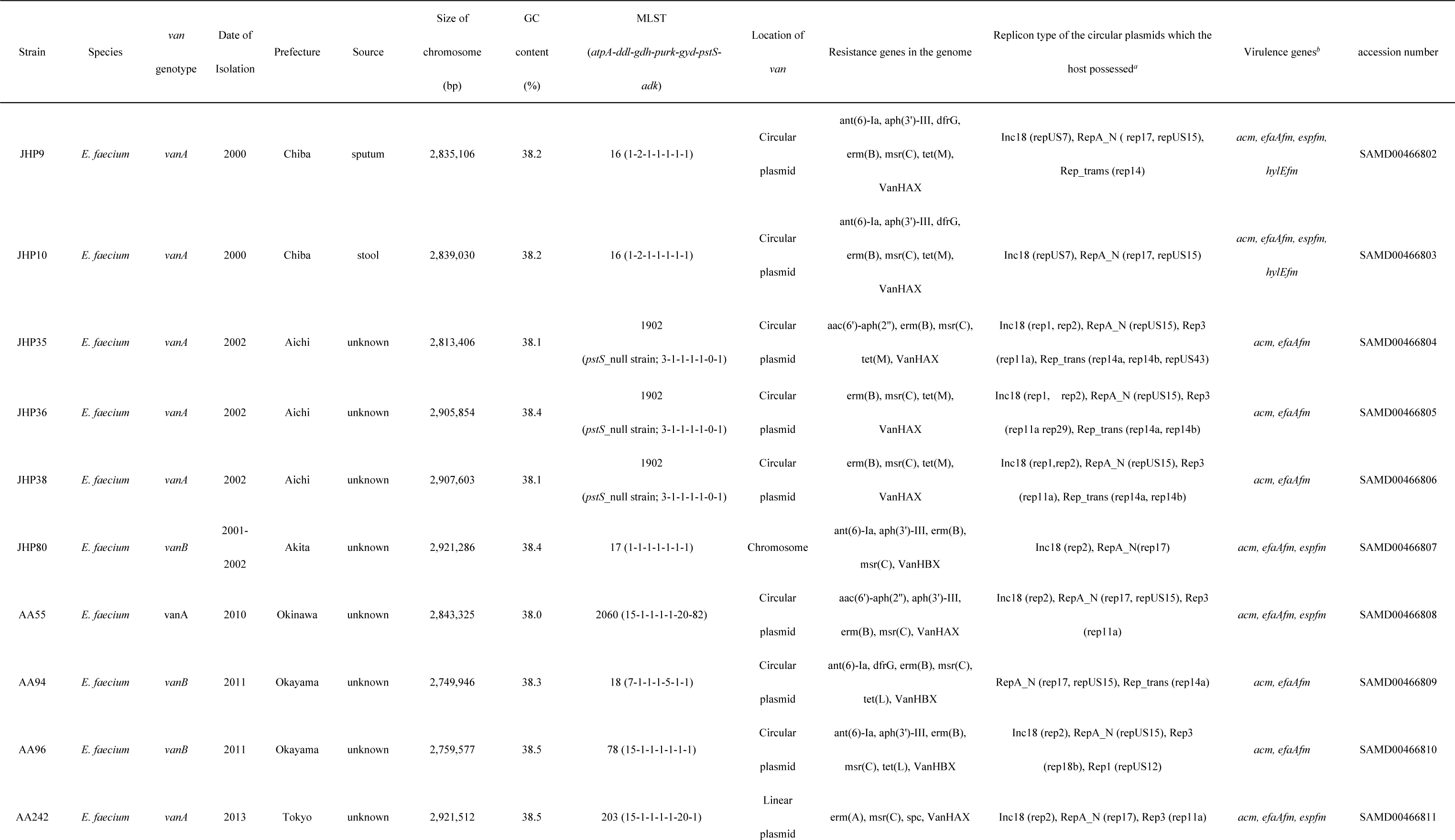
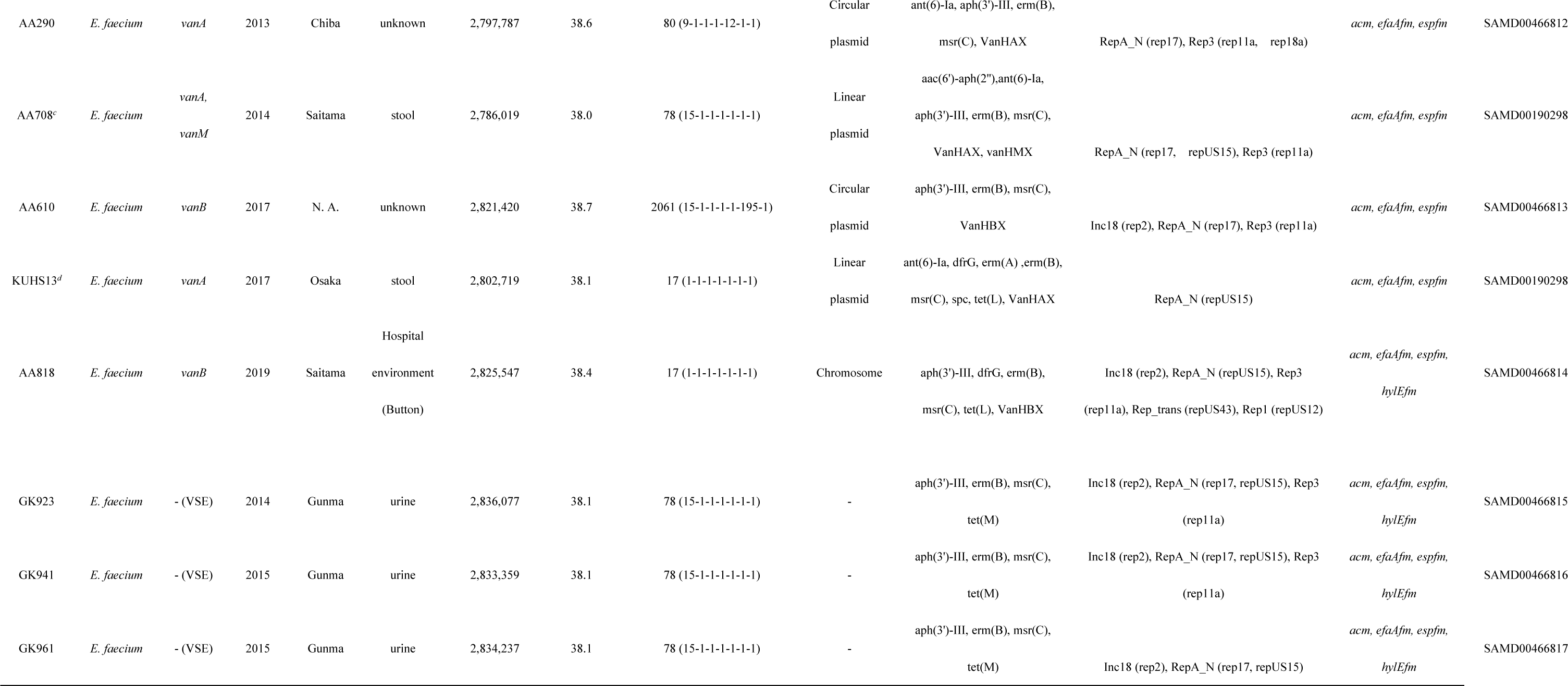
Strains carrying pELF1-like plasmids. N.A. stands for not available. *^a^*The replicon type was examined by default settings using the PlasmidFinder (2.0.1). *^b^*The virulence genes were identified using the Virulence Finder (2.0). *^c, d^*Strains that have already been reported previously.

Using WGS with multi-locus sequence typing (MLST), we found that, of the 16 strains, two were ST16, two were ST17, one was ST18, one was ST78, one was ST80, one was ST203, and three were *pstS*-null strains (ST1902). The nucleotide sequences for *adk* of AA55 and *pstS* of AA610 did not match the previously reported alleles and showed new ST numbers (ST2060, and ST2061, respectively). The three VSE strains were ST78. All 16 strains belonged to clonal complex 17 (CC17), an important factor of nosocomial infection^3, 10^.

Except for strain JHP35, all VanA-type VREs were resistant to both vancomycin (>512 mg/L) and teicoplanin, whereas all VanB-type VREs were relatively less resistant to vancomycin (32–64 mg/L) and susceptible to teicoplanin, preserving the characteristics of each resistant type (Supplementary Table 2). All strains were resistant to ampicillin, which is consistent with the characteristics of CC17^3^.

According to the PCR results, the presence of the pELF1-like plasmid was confirmed in all 16 strains (Table 3). Each pELF1-like plasmid was named using the strain name (e.g., pELF_JHP9). We examined the replicon of the plasmids using Plasmidfinder (Table 2)^11^, and found that all the strains contained one or more circular plasmids, suggesting that pELF1-like plasmids coexist with various circular plasmids belonging to the Inc18-, RepA_N-, Rep3-, and rolling-circle replicating (RCR)-type (Rep_trans and Rep1) plasmids (Table 2). Regarding virulence genes, all strains had *efaAfm* and *acm* on their chromosomes, but none were identified on the pELF1-like plasmids (Table 2)^12–14^.

**Table 3.**
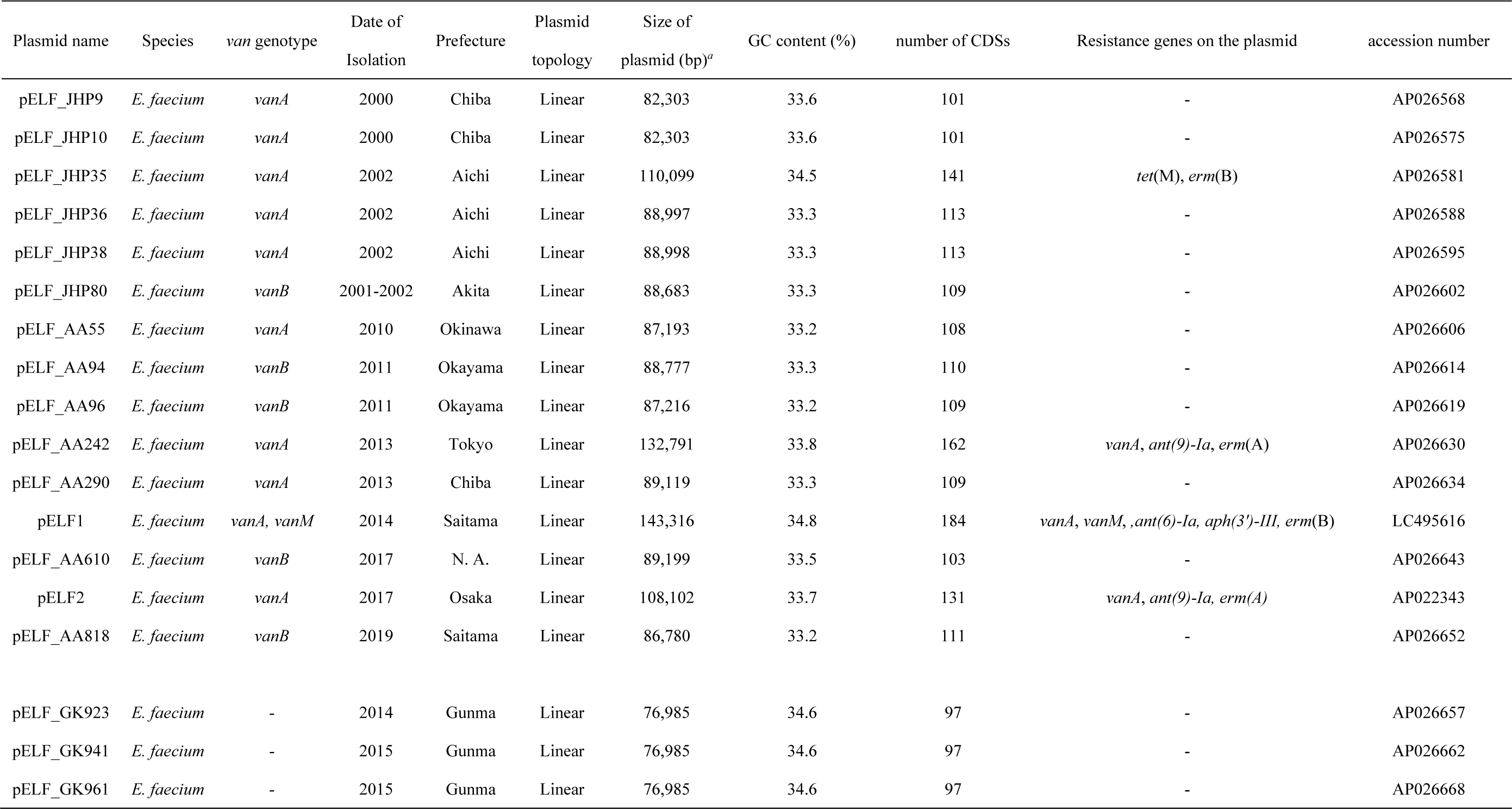
The pELF1-like plasmids found in the Japanese clinical isolates. N. A.; not applicable

### Comparison of pELF1-like plasmid genetic structures based on WGS

pELF1-like plasmids are unique in that their opposing ends have different structures (known as a hybrid-type linear plasmid)^7^. We investigated the structure of the terminal ends of all pELF1-like plasmids using WGS and found that the nucleotide sequences at both ends were highly conserved and presumed to have different structures, consistent with pELF1 (Supplementary Fig. 2).

Few reports describe the diversity of the genetic structure of pELF1-like plasmids. Accordingly, we conducted a search of the NCBI database based on the whole nucleotide sequence of pELF2. We selected plasmids with linear topology from enterococci as candidates and identified 14 pELF1-like plasmids with conserved terminal ends (Table 4) from a variety of countries, including Denmark, Norway, China, the United States, Switzerland, India, and the Netherlands (Supplementary Fig. 1B). The hosts of these 14 pELF1-like plasmids were all *E. faecium*, and all belonged to CC17.

**Table 4.**
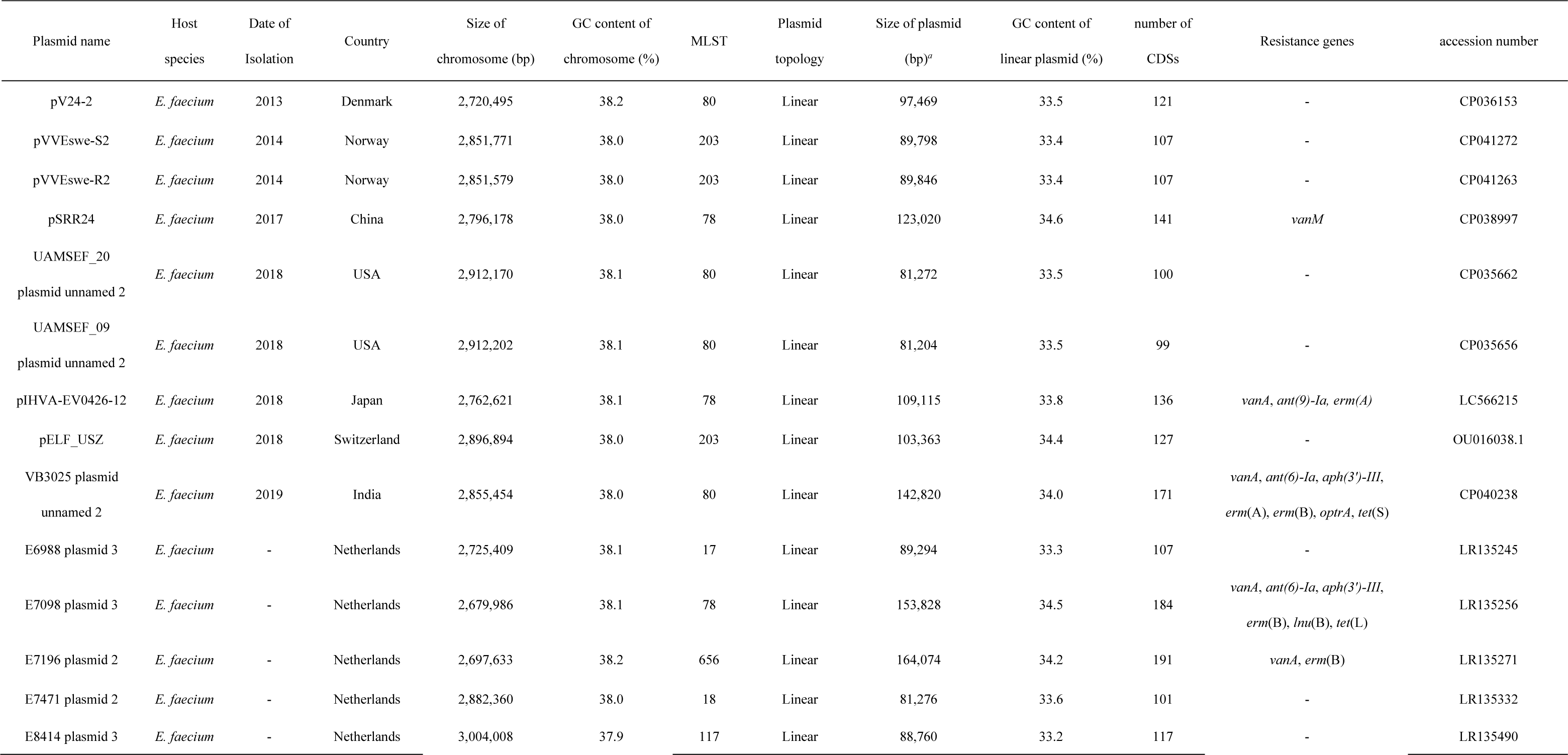
The putative pELF1-like plasmids and their hosts found in the public database. Plasmids with presumed linear topology were retrieved from the NCBI database. *^a^*The nucleotide sequence sizes of the plasmids with both ends modified with reference to pELF1 are described.

pELF1-like plasmid size was relatively diverse, with the shortest length being 76 kb for pELF_GK923, GK941, and GK961 and the longest being 164 kb for E7196 (Tables 3 and 4). The minimum number of coding sequences (CDS) was 97 for pELF_GK923, GK941, and GK961, and the maximum number was 191 for E7196. We also found that the GC content of the pELF1-like plasmid was ∼4% lower than that of the host genome (mean GC content: 33.9% vs. 38.1%; Tables 2–4). Furthermore, pELF1-like plasmids exhibited lower GC content than circular plasmids harbored by multiple *Enterococcus* spp., including *E. faecium* (Mann–Whitney test (*p* < 0.01); Supplementary Fig. 3, and Supplementary Table 3).

We performed a core genome analysis with 18 plasmids, including pELF1 and pELF2, along with 14 putative pELF1-like plasmids confirmed by a public database search (Fig. 1 and Supplementary Table 4)^9, 15^. At Blastp = 80% (considered to cover most core genes), a rapid large-scale prokaryote pan-genome analysis (Roary) estimated the core genome to consist of 57 CDSs, which was 56.4% of the CDSs in pELF_JHP9 (Supplementary Table 4). According to the annotation data, most of the 57 core genes were hypothetical proteins (Supplementary Table 5). Among the core genes, we identified some encoding replication proteins and several toxin–antitoxin (TA) system-related genes.

**Fig. 1.**
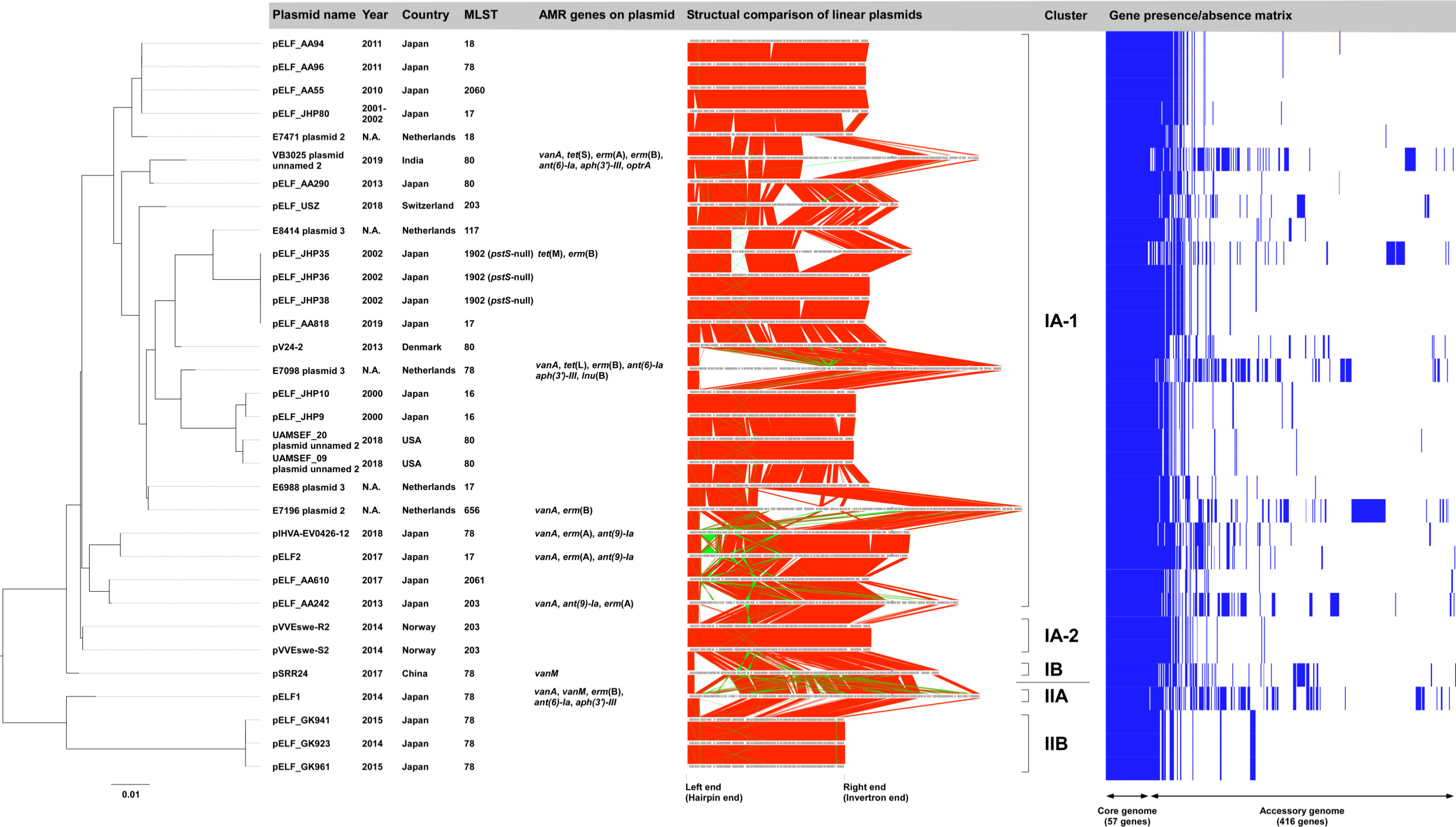
Comprehensive phylogenetic analysis of pELF1-like plasmids. Core genome analysis was performed using Roary, and phylogenetic trees were generated in RAxML. Plasmid names, isolation years, countries, MLST, and AMR gene possession status, and representative figures are shown along with the phylogenetic tree results. Synteny block lineages in the same orientation are shown in red, while those in the reverse orientation are shown in green. The gene presence/absence matrix was constructed using Phandango.

We performed a phylogenetic analysis based on the core gene sequences and identified several genetically distinct plasmid clusters (Fig. 1), which could be classified into two major groups—Cluster I and Cluster II. Cluster I was classified into IA and IB, and Cluster II into IIA and IIB. Cluster IA was further classified into IA-1 and IA-2. Cluster I accounted for the majority of plasmids, among which Cluster IA-1 comprised 25 of the 32 pELF1-like plasmids. Cluster II contained four pELF1-like plasmids, including pELF1, pELF_GK923, 941, and 961, which were harbored by three VSE strains. Cluster IA-1 accounted for pELF1-like plasmids detected worldwide, while Cluster IA-2 accounted for the plasmids from Norway, Cluster IB those from China, and Cluster II those from Japan. pELF_AA94 and pELF_AA96 (harbored by strains detected in the same year in the same prefecture in Japan) belonged to the same cluster, and their synteny block lineage suggested a high degree of similarity (pairwise nucleotide identity = 98.2%); however, their host MLST types were different. This finding suggests the occurrence of plasmid transfer in clinical settings. Similarly, pELF_JHP36 and pELF_AA818 were highly similar plasmids (pairwise nucleotide identity = 97.5%), with different host MLST types. Their detection was 17 years apart, and they were detected in different regions of Japan. Likewise, the pELF_JHP9/pELF_JHP10 from Japan and UAMSEF_20 plasmid unnamed 2/UAMSEF_09 plasmid unnamed 2 from the US belonged to the same cluster (pairwise nucleotide identity = 90.8%). These findings suggest that pELF1-like plasmids are structurally stable and occur worldwide.

We also found that the pELF1-like plasmids detected in Japan were clustered into different lineages (Fig. 1), though with highly conserved genetic structures. We could not confirm any major genomic rearrangement, such as inversion, which would change the order of core genes (Fig. 1). Most of the synteny blocks displayed in reverse orientation were insertion sequences (ISs), with the only exception being the reverse orientation of the antimicrobial resistance (AMR) region containing the *vanA* gene cluster between pELF2 and pIHVA-EV0426-12 (Supplementary Fig. 4). While the pELF1-like plasmid backbone was highly conserved, its diversity resulted from MGEs (Figs. 1 and 2).

**Fig. 2.**
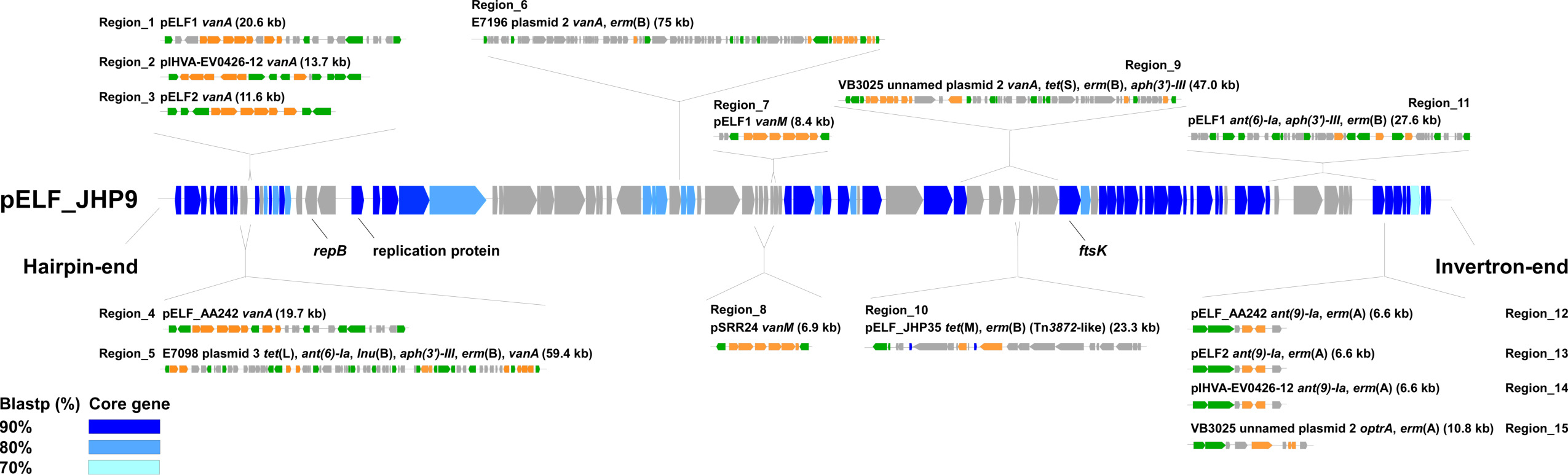
Genetic structure of the core genes and AMR regions of pELF1-like plasmids. The locations of the core genes of the 32 pELF1-like plasmids and the insertion sites of the AMR regions of each pELF1-like plasmid were described in pELF_JHP9. The core genes obtained for each BLASTP cut-off value are shown in blue gradient panels. For the AMR region, the plasmid name, drug resistance gene name, size (kb), and genetic structure schemas are noted along with the insertion position. The gray panel shows non-core genes. The green panels represent transposition/site-specific recombination-related genes, and the orange panels represent AMR genes.

### Analysis of the AMR region generating the diversity of pELF1-like plasmids

Next, we focused our analysis on the AMR region comprising the accessory genome. A Resfinder search showed that 9 of 32 pELF1-like plasmids harbor AMR genes (Tables 3 and 4). Since the order of CDSs in the pELF1-like plasmids was extremely well preserved, the insertion sites of AMR regions were assessed based on pELF_JHP9, one of the shortest plasmids belonging to Cluster IA-1 (Figs. 1 and 2; Supplementary Figs. 5 and 6).

We identified 15 AMR regions in 32 pELF1-like plasmids. Nine of the AMR regions contained *van* gene clusters with IS*1216E* on at least one of the terminal ends, indicating that the IS*1216E*-translocatable unit is involved in AMR transmission onto pELF1-like plasmids (Region_1–9, Supplementary Fig. 5). Five of the AMR regions had IS*1216E* at both ends, four in the same orientation and one in the opposite orientation (Region_1, 4–6, 9; Supplementary Fig. 5). Five of the AMR regions containing the *vanA* gene cluster were concentrated to the region near the left hairpin end, and the AMR region containing the *vanM* gene cluster was concentrated to the center of the pELF1-like plasmid (Region_7, 8; Fig. 2 and Supplementary Fig. 5). Similar to the AMR region containing the *van* gene cluster, IS*1216E* was also present at the extremities of the AMR region containing *ant(6)-Ia*, *aph(3’)-III*, and *erm*(B) of pELF1 (Region_11; Fig. 2 and Supplementary Fig. 6). pELF2 and pIHVA-EV0426-12 possessed an AMR region similar to those containing *ant(9)-Ia* and *erm*(A) of pELF_AA242 (Region_12–14; Fig. 2 and Supplementary Fig. 6). VB3025 plasmid unnamed 2 possessed a similar AMR region, but with *optrA* instead of *ant(9)-Ia* (Region 15; Fig. 2 and Supplementary Fig. 6). These four AMR regions were located near the core gene at the invertron-end of the plasmid (Fig. 2). In these AMR regions, the presence of XerC1 and -C2 suggested their involvement in transitions (Supplementary Fig. 6). pELF_JHP35 harbored Tn*916*, whose *orf9* was disrupted by Tn*917*. This closely resembled the Tn*3872*-like element of CGSP14 (*Streptococcus pneumoniae*, Region 10; Supplementary Figs. 6 and 7)^17^.

### Phenotypic analysis of fitness cost and stability of pELF1-like plasmids in enterococcal hosts

Plasmids allow for the development of environmentally adaptive traits, such as drug resistance genes, but they also impose fitness costs^18^. Fitness cost is an important factor of long-term stability in plasmids, especially under conditions where the genes encoding for drug resistance are redundant^19, 20^. We analyzed the fitness cost and stability of pELF1-like plasmids using pELF2, which harbors several AMR genes, including *vanA*. This pELF1-like plasmid belongs to Cluster IA-1 and is known to be spreading in some regions of Japan (Fig. 1)^8^.

To investigate host-specific differences, we tested four strains under non-selective conditions at temperatures mimicking that of the human body (37 °C), environmental wastewater (25 °C), or chicken body (42 °C, Fig. 3a and 3b)^21–23^. In *E. faecium*, there was no significant reduction in the growth curve related to pELF2 at any temperature. In contrast, a clear decrease in the growth curve of *E. faecalis* was observed at all temperatures. pELF2 imposed a fitness cost on all four enterococci species; however, the fitness costs for *E. faecium* and *E. casseliflavus* were the lowest, with 4.1% and 3.2%, respectively, at 37 °C (Supplementary Table 6). For these two species, the fitness cost did not increase at any of the temperatures (Fig. 3b). In contrast, we observed several times higher fitness costs for *E. faecalis* and *E. hirae*. To confirm whether this physiological effect was pELF2-specific, experiments with another pELF1-like plasmid (pELF2_AA242) were performed, which confirmed similar growth effects and fitness costs (Supplementary Fig. 8a and 8b and Supplementary Table 6). For *E. faecalis*—the most prevalent among human enterococci infections—we changed the host strain to OG1RF (accession number: NZ_ABPI00000000.1) and analyzed the growth effect of the pELF1-like plasmids (Supplementary Fig. 8C and 8D). As with the FA2-2 (*E. faecalis*) host, we observed significantly lower growth rates and higher fitness costs, which implies that the fitness cost imposed by the pELF1-like plasmid involves host-specific factors. Accordingly, we assessed codon usage frequency; the codon adaptation index (CAI) between host genomes and MGE is hypothesized to correspond with inhibition of gene translation and negatively impact fitness costs^24^. In *E. faecium*, there was no significant difference in CAI between genes on the chromosomal genome and pELF2. The difference was greatest in *E. faecalis* (median CAI of *E. faecalis* chromosomal genes vs. pELF2 genes; 0.5090 vs. 0.4690, *p* < 0.0001 [Mann–Whitney test]; Supplementary Fig. 9).

**Fig. 3.**
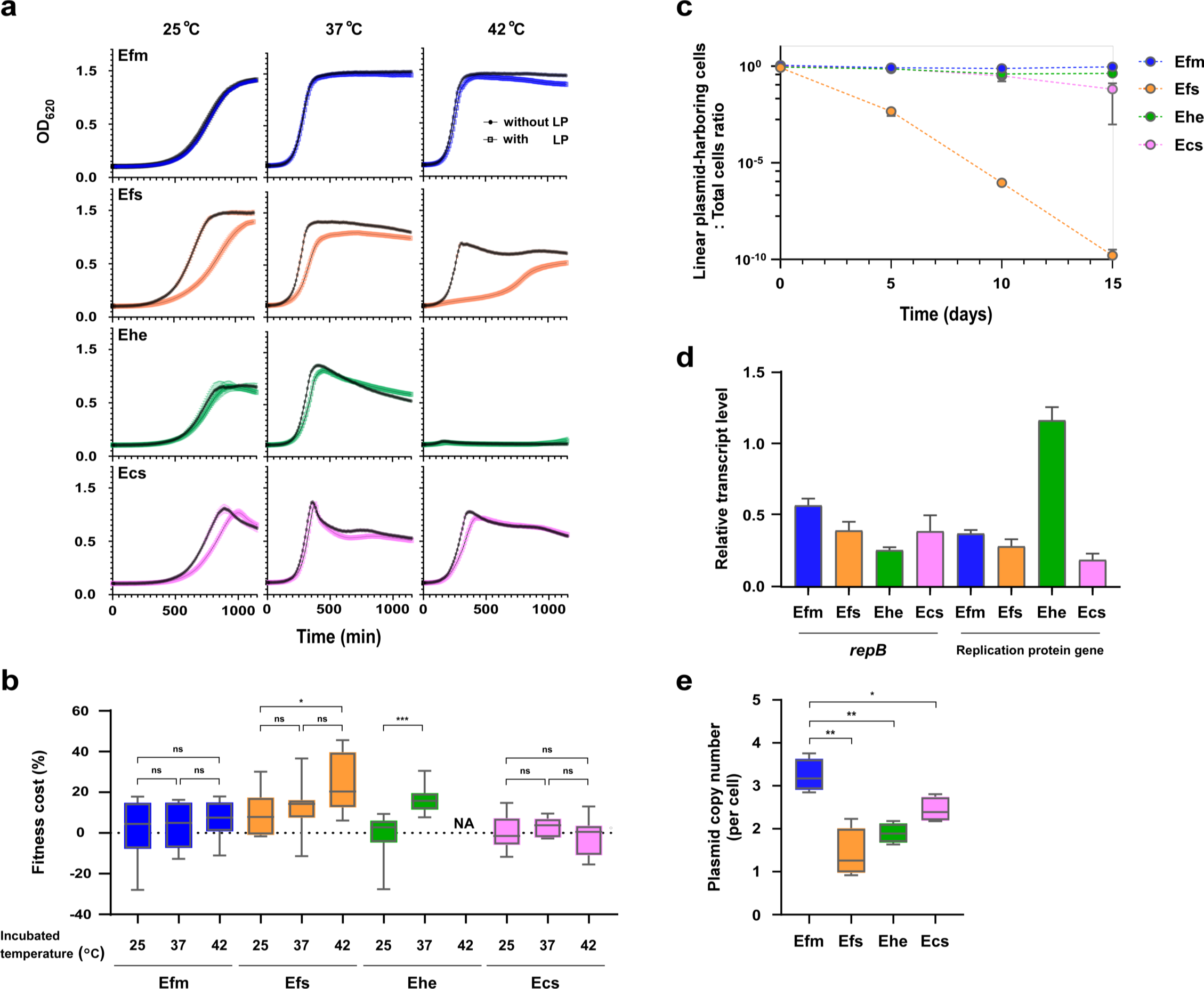
Analysis of the stability and effect of the pELF1-like plasmid on the growth of enterococci. The effects of pELF1-like plasmid (pELF2) carriage on the (a) growth curve and (b) plasmid-imposed fitness cost on four *Enterococcus* spp. are shown. For the analysis of growth and fitness cost, nine biological replicates of the experiment were conducted. (a) Growth curves are shown as absorbance (OD_620_) ± SE. Black circles indicate pELF2 non-carrying strains and white squares indicate pELF2-carrying strains. (b) Fitness cost is shown in a box-and-whisker diagram. The whiskers indicate the minimum to maximum values. *, *p* ≤0.05; ***, *p* ≤0.001; ns, not significant; NA, not applicable (Mann–Whitney test). (c) The stability of the pELF1-like plasmid was evaluated in a 15-d passaging experiment in a liquid medium without selection pressure. After 5, 10, and 15 d, the number of pELF1-like plasmid-harboring and non-harboring strains was measured, and the ratio (± SE) of the plasmid-harboring cells to total cells was calculated. Three independent biological experiments were conducted. (d) The transcript level of *repB* and replication protein genes were measured using reverse transcription (RT)-qPCR. The bar charts show the relative transcript ratio compared to mRNA levels of each *gyrB* gene, along with ±SE. (e) Plasmid copy number was measured using qPCR and shown in a box-and-whisker diagram. *, *p* ≤0.05; **, *p* ≤0.01 (unpaired *t*-test; *p* = 0.0019, *p* = 0.0010, and *p* = 0.0176, respectively). Blue symbols indicate *E. faecium* (Efm, BM4105RF), orange symbols indicate *E. faecalis* (Efs, FA2-2), green symbols indicate *E. hirae* (Ehe, ATCC9790RF), and pink symbols indicate *E. casseliflavus* (Ecs, KT06RF).

We assessed the stability of pELF1-like plasmids in four *Enterococcus* spp. via long-term passage assay under non-selective conditions. During 15-day passages, pELF2 maintained the highest stability in *E. faecium*, followed by *E. hirae* and *E. casseliflavus* (Fig. 3c). However, in *E. faecalis*, the percentage of pELF1-like plasmid-carrying strains markedly decreased as passaging progressed, falling below the detection limit at 30 passages (15 d). To confirm whether fitness improved within a declining pELF2-carrying *E. faecalis* population, we investigated the genetic structure of pELF2 and fitness cost in *E. faecalis* after 5 d of passage (day 5-evolved strains). Pulsed-field gel electrophoresis (PFGE) showed a marked reduction in the size of pELF2 in some of the day 5-evolved strains (1.1 and 1.2 strains) (Supplementary Fig. 10a), which also implies that pELF2 is unstable in *E. faecalis*. Somewhat surprisingly, given the eventual disappearance of pELF2 from the *E. faecalis* population, day 5-evolved strains had a 20–30% reduction in fitness cost compared to parental strains (Supplementary Fig. 10b). Furthermore, the plasmid copy number of these day 5-evolved strains was higher than that of the parent strain (Supplementary Fig. 10c). We tested the plasmid transfer frequencies possessed by these day 5-evolved strains and found that the plasmid of 1.1 strain, which showed a decrease in plasmid size, had lost self-transmissibility (Supplementary Fig. 10d). SNP analysis based on WGS data of the 1.1 strain did not reveal any mutations in the coding sequence of pELF2 (Supplementary Table 7); however, mapping analysis identified that 1.1 strain-harboring pELF2-derivative had lost a 31.6 kb region containing several core genes (Supplementary Fig. 10e).

Since overexpression of replication-associated proteins is known to impair genome replication and lead to fitness costs, we examined the transcription levels of *repB* and the replication protein gene on pELF2 in four different hosts. We found that the transcription levels of *repB* and the replication protein gene were lower than those of *gyrB* on each chromosome (Fig. 3d)^25^. Using qPCR, we tested whether the copy number of pELF1-like plasmids varied between *Enterococcus* spp. (Fig. 3e and Supplementary Table 8). Consistent with the RT-qPCR results, the plasmid copy number appeared to be tightly controlled in all four *Enterococcus* spp., as with other conjugative plasmids^26^. However, we observed interspecific differences in the number of plasmid copies, with *E. faecium* possessing significantly more pELF1-like plasmid copies (3.2 copies/cell) than the other three species. The lowest number of plasmid copies was observed in *E. faecalis*, which has the highest pELF2 fitness cost.

### The impact of pELF1-like plasmids on *E. faecium*

Plasmid carriage impacts bacterial physiology, e.g., metabolic regulation, but the effects of pELF1-like plasmids on gene transcription are unclear^27, 28^. To analyze the interaction between the pELF1-like plasmid and the host chromosome, we compared the levels of chromosomal gene transcription between pELF2-carrying *E. faecium* and non-carrying *E. faecium*^29^. Of the 2,515 genes, 56 genes (∼2%) showed a >2-fold change in the transcript level, including 38 upregulated and 18 downregulated genes (Adjusted *p*-value < 0.05; Fig. 4a and Supplementary Table 9), indicating a modest effect of pELF1-like plasmid carriage^30^. Among the upregulated genes, we identified glutamine-fructose 6 phosphate aminotransferase (*glmS_1*) and glutamine synthase (*glnA*). These genes are involved in the synthesis of glucosamine 6 phosphate, which participates in UDP-NAcGlc synthesis and metabolism of the cell wall structure. Of the downregulated genes, 50% were related to carbohydrate transport and metabolism (Supplementary Table 9). A gene ontology (GO) analysis showed that carbohydrate transport and the phosphoenolpyruvate-dependent sugar phosphotransferase system (PTS, especially mannose-specific PTS) were significantly downregulated in the clusters of orthologous group (Fig. 4b)^31^. Similar to the transcriptome analysis of *K. pneumoniae* plasmid acquisition, pELF2 carriage influenced metabolism-related genes^28^. To investigate its effect on carbohydrate availability, we conducted growth experiments under carbohydrate-limited conditions. Contrary to the RNA-seq results, we did not observe any change in growth in the presence of glucose or mannose with or without the pELF1-like plasmid (Supplementary Fig. 11).

**Fig. 4.**
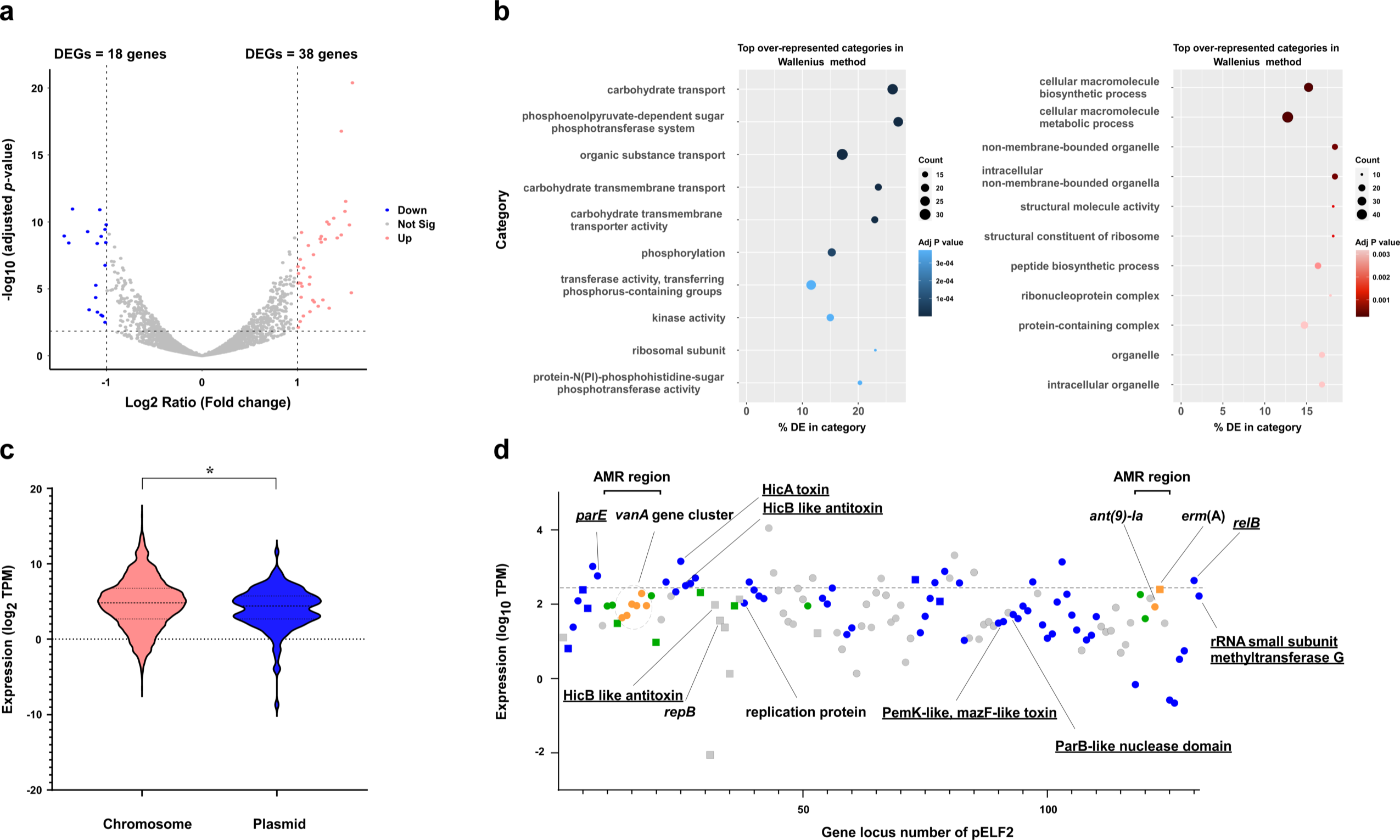
Transcriptome analysis of the effect of pELF1-like plasmid carriage on *E. faecium* host genome. We assessed the effect of pELF2 carriage on the *E. faecium* host (BM4105RF) via transcriptomic analysis. (a) Changes in the transcriptome of *E. faecium* possessing pELF2. Each dot indicates an open reading frame, with red circles indicating upregulation (log_2_ fold change > 1, adjusted *p*-value < 0.05) and blue circles indicating downregulation (log_2_ fold change < –1, adjusted *p*-value < 0.05). (b) Gene enrichment analysis was performed with GOseq based on DESeq2 results. The blue and red bubble charts show the results of the downregulated and upregulated gene analysis. (c) Transcript abundances of genes on the chromosome (blue) and the pELF1-like plasmid (red, log_2_ of transcript per million, TPM). The transcript abundances of the genes on the plasmid were calibrated with the copy number of the plasmid. *, *p* < 0.05 (Kolmogorov–Smirnov test). (d) The TPM values of CDS of pELF2 are plotted in the order of gene locus number. The circles indicate the plus strand and the squares indicate the minus strand. Dark blue indicates core genes; among non-core genes, IS and recombinase are shown in green, drug resistance genes are shown in orange, and other genes are shown in gray. The dashed line indicates the Top 20% TPM values.

### Transcriptome analysis of pELF1-like plasmid genes in *E. faecium*

We assessed the transcriptional profile of genes on pELF2 and found that the transcript level of genes on the pELF1-like plasmid was lower than that on the chromosomes (*p* = 0.01 [Kolmogorov–Smirnov test], Fig. 4c). This finding is consistent with the characteristics of the conjugative plasmid of *P. aeruginosa*, which is associated with reduced fitness costs and presumably promotes the stability of the pELF1-like plasmid^18, 26, 32^.

pELF2 is a natural plasmid that has spread in some areas of Japan and possesses multiple drug resistance genes^8^. We compared the transcript levels of the core genes, non-core genes, and genes on MGEs (including AMR genes), but found no significant differences (Supplementary Fig. 12). The low transcription levels of the entire pELF2 gene set implicated the regulation of the transcription of relatively expensive core genes, such as transfer-related genes, while also suggested that the accessory genes in pELF2 do not impose a high cost to *E. faecium*.

Of the genes on pELF2 in the top 20% of transcript per million (TPM) values, 65% were core genes, and four genes encoding the TA system were identified (Fig. 4d and Supplementary Table 5). The TA system is considered to contribute to robust vertical propagation by post-segregational killing (PSK), which is particularly important in low-copy plasmids^33^. According to TASer and BLAST searches, multiple putative Type II TA system-related genes are present in pELF2, of which 5 were core genes^34^. The Type II TA system is the most abundant type in bacterial genomes, including MGEs^35^. HicAB was classified as a Type IIA TA system (harbored by many bacteria) and transcriptome analysis showed high transcription levels^36^. Similarly, the transcript levels of the RelE/ParE family toxin, and RelB/DinJ family antitoxin were comparably high in the top 20% of TPM values^37^. From these results, we hypothesized that these relatively highly transcribed TA systems contributed to the stable inheritance of pELF1-like plasmids in *E. faecium*^38–40^.

### Comparative analysis of pELF1-like plasmid-carrying *E. faecium* host genomes

The above analysis of epidemiological data and physiological effects suggest that *E. faecium* is an optimal host. However, the stability of plasmids varies even within the same bacterial species with strain-specific traits^41^. To determine the genetic background of the pELF1-like plasmid carrying *E. faecium*, we conducted a phylogenetic analysis of the complete genome of 309 *E. faecium* strains, including 32 pELF1-like plasmid-carrying strains (Fig. 5 and Supplementary Table 10). The phylogenetic tree was divided into two major groups, with the smaller group containing strains of Clade B^42, 43^; therefore, these two groups were considered to correspond to the previously reported clade classification. The pELF1-like plasmid-harboring strains all belonged to CC17, and, being ampicillin-resistant, were considered to belong to Clade A1^3, 43^. Even within Clade A, they were well-dispersed and not concentrated to any particular lineage. These results indicate that the pELF1-like plasmid could be carried by various lineages of *E. faecium* belonging to the MDR clade.

**Fig. 5.**
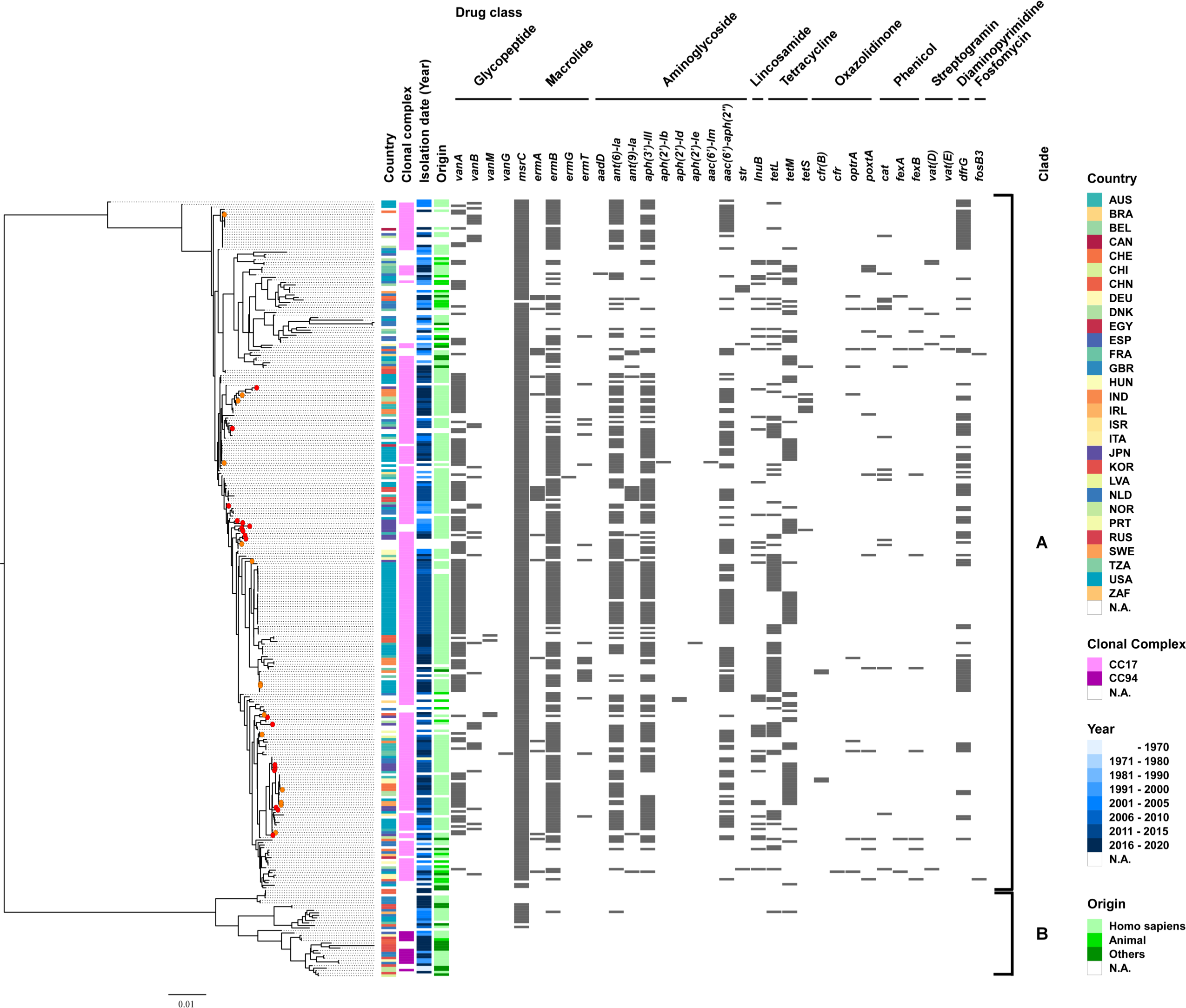
Genomic comparison of pELF1-like plasmid-harboring vs. non-harboring *E. faecium* strains based on the core genome. Phylogenetic tree analysis based on core genome SNPs was performed on 309 strains of *E. faecium*, including 32 strains harboring pELF1-like plasmids. Core genes were identified using the Roary, with a cut-off value of Blastp = 95%. The panel shows the country, clonal complex, year, and origin of each strain, as well as the resistance gene status. For the clonal complex, CC17 and CC94 were described based on the MLST profile in PubMLST (https://pubmlst.org/organisms/enterococcus-faecium). The clade was determined based on the phylogenetic tree of Lebreton *et al.*.^43^ The red circles in the phylogenetic tree indicate pELF1-like plasmid-harboring strains of Japanese origin, and the orange circles indicate pELF1-like plasmid-harboring strains in the public database.

## Discussion

MGEs, such as plasmids, play an important role in the efficient propagation of drug resistance genes in enterococci. Several reports on enterococcal pELF1-like plasmids have shown that they confer vancomycin resistance and promote carbohydrate availability; although they have a significant impact in clinical practice, their biological effects on the host enterococci are poorly understood^8, 9, 15^. In this study, we conducted a comprehensive analysis of pELF1-like plasmids based on growth phenotype and transcriptomics coupled with molecular epidemiology.

We found that 3.4% of VREs and 0.2% of VSEs harbored pELF1-like plasmids, and all were *E. faecium* (Table 1). As early as the beginning of 2000 in Japan, pELF1-like plasmids were detected in vancomycin-resistant *E. faecium* (Table 2). In combination with the results of public database searches, phylogenetic analysis based on the core genome of the 32 pELF1-like plasmids showed that 1) multiple lineages with a high degree of structural similarity exist and 2) Cluster I pELF1-like plasmids occur globally (Fig. 1). The genetic structure and order of CDSs of the pELF1-like plasmids, detected over ∼20 years, were highly conserved (Fig. 1). The pELF1-like plasmid also had the plasticity to acquire accessory genes (Fig. 2). The MGEs containing the AMR genes were transferred to several pELF1-like plasmids (Fig. 2). Of the 15 AMR regions on the pELF1-like plasmid, IS*1216E* was present at a comparatively high frequency (10 of 15 AMR regions; Fig. 2 and Supplementary Fig. 5 and 6). The IS*6* family, to which IS*1216* belongs, is involved in the propagation of AMR genes, both gram-positive and -negative, and is reportedly responsible for many transitions between plasmids and chromosomes^40, 44^. According to the MobileElementFinder-based search, pELF1-like plasmids with AMR genes retained more MGEs, including ISs (AMR-holding plasmid vs. non-holding plasmid = 7.25 vs. 0.64 [Mean number of MGEs per pELF]; Supplementary Table 11)^45^. These findings revealed that IS-associated AMR regions are primarily responsible for the diversity of pELF1-like plasmids.

To confirm their adaptation to enterococci, especially *E. faecium*, we investigated the phenotypic effects of pELF1-like plasmids on *Enterococcus* spp. growth. The persistence of a conjugative plasmid in a bacterial population is believed to be determined by 1) the frequency of conjugative transmission, 2) fitness cost imposed by the plasmid, and 3) loss of the plasmid during cell segregation^19, 20^. Therefore, both horizontal transmission and vertical inheritance are involved in the long-term plasmid stability. The pELF1-like plasmids exhibit a high transfer frequency in both solid and liquid mediums, which contributes to their persistence^7, 8^. In addition, the fitness cost of the plasmid must be sufficiently low for stable vertical inheritance^18^. As with the results of the plasmid stability experiments, we observed a lower impact on growth and fitness cost related to pELF1-like plasmids in *E. faecium* under all temperature conditions (Fig. 3a–3c, and Supplementary Table 6). We observed similar trends in fitness cost for the other pELF1-like plasmids (pELF_AA242), suggesting that their effect on enterococci is a general phenomenon (Supplementary Fig. 8).

Various origins of fitness cost have been suggested, one of which is the disruption of genome replication resulting from overexpression of replication-associated genes^18, 25^. Plasmid copy number is also considered to increase fitness cost, but the pELF1-like plasmid had no noticeable effect on the transcription of replication-related proteins and its copy number was relatively low (Fig. 3d and 3e). This result was observed not only for *E. faecium*, but also for *E. faecalis*, which showed a high fitness cost associated with pELF2. Since most of the fitness costs are considered to arise from the expression of genes on the plasmid, the effective control of transcription is crucial^18^. Transcriptome analysis revealed that transcription of entire genes on the pELF2 plasmid was suppressed to a lower level than in the chromosomal genome, and the effect of plasmid carriage on host chromosomal genes was limited (Fig. 4). Together with the lack of significant differences in CAI, these results could explain the lower fitness cost in *E. faecium*.

Meanwhile, in *E. faecalis*, where pELF2 imposes a high fitness cost, the transfer-related region could be one of the factors in the cause of the fitness cost (Supplementary Fig. 10). This result was consistent with reports that deletion of the region containing the conjugation machinery reduced plasmid fitness cost^49^. In the day5-evolved strain (1.1 strain), the fitness cost of the pELF2-dertivative was reduced; however, the change do not appear to be sufficient for the establishment of the plasmids in the *E. faecalis* population (Fig. 3c and Supplementary Fig. 10). This suggested that plasmid transfer also had a critical role in the persistent of the plasmids.

Given the influence of pELF1-like plasmid genes on plasmid stabilization, we hypothesized that a large number of highly transcribed TA systems promote stable vertical inheritance (Fig. 4d and Supplementary Table 5), although it may also be related to the expansion of the host range of plasmids, as previously reported ^46^. Consistent with previous reports of nosocomial transmission, we confirmed that the pELF1-like plasmid was stably maintained by multiple *Enterococcus* spp.^8^. The wide range of stable hosts would likely promote plasmid diversity and contribute to the long-term survival of plasmids in a variety of settings, including clinical practice^47^. Thus, the pELF1-like plasmid is a low-cost plasmid (especially for *E. faecium*), with high stability, high frequency of self-transmissibility, and the ability to carry AMR genes, which makes it an ideal candidate for spreading drug resistance between *Enterococcus* spp.

The core genome analysis revealed that various lineages of *E. faecium* belonging to the MDR clade carry pELF1-like plasmids (Clade A1, Fig. 5). It is worth noting that, of the 32 pELF1-like plasmids in this analysis, all pELF1-like plasmids with vancomycin resistance genes were detected after 2010. More recently, Egan *et al.*^48^ reported evidence of the regional spread of pELF1-like plasmids carrying the *vanA* gene cluster in Ireland. These findings indicate that pELF1-like plasmids have only recently become important vehicles for vancomycin resistance genes, at least in some regions^8, 9^.

In this study, we combined molecular epidemiological, phenotypic, and transcriptomic analyses to provide an integrated characterization of the pELF1-like plasmid. Our characterization of the relationship between pELF1-like plasmids and *E. faecium*, the most abundant VRE, highlights the importance of further analysis. We acknowledge some limitations to our research. Our in silico and in vitro analyses neglected the context-specific effects of plasmids *in vivo*, though this was unavoidable given that the adaptation of pELF1-like plasmids to enterococci *in vivo* remains unclear. In addition, many of the strains used were Clade A medical strains, with Clade B or enterococci of environmental origin being underrepresented. Future studies should also consider the spread of pELF1-like plasmids in animals and food from a one-health perspective.

## Materials and Methods

### *Enterococci* strains and drug susceptibility test

The bacterial strains used in this study were stored at the Department of Bacteriology, Gunma University Graduate School of Medicine, Gunma, Japan (Tables 1 and 2). All strains were detected at medical institutions in Japan. Unless otherwise noted, enterococci were cultured in Todd–Hewitt broth (THB; Difco Laboratories, Detroit, MI, USA) at 37 °C in a static condition. Minimum inhibitory concentrations (MIC) were determined using the agar dilution method, and interpretation of MIC results was based on the Clinical and Laboratory Standards Institute guidelines (http://clsi.org/). NCBI was used to search for and retrieve genomes^49^. Transconjugants were obtained via filter mating experiments as previously reported^7^. Vancomycin (5 mg/L) or erythromycin (16 mg/L) was used for transconjugant selection along with both rifampicin (50 mg/L) and fusidic acid (50 mg/L). KUHS13 (accession number: SAMD00202474) and AA242 (accession number) were used as donor strains, and FA2-2 (*E. faecalis*; accession number: SAMN00809127), BM4105RF (*E. faecium*; accession number: SAMN09464428), ATCC9790RF (*E. hirae*; accession number: SAMN02604142), and KT06RF (*E. casseliflavus*) were used as recipient strains. In the second conjugation experiment, we used BM4105SS (*E. faecium*) as a recipient strain. Vancomycin (5 mg/L) was used for transconjugant selection along with both streptomycin (516 mg/L) and spectinomycin (256 mg/L). Before experimentation, we confirmed the presence of pELF1-like plasmids using PCR and PFGE.

### Colony PCR

We performed colony PCR to detect pELF1-like plasmids. Briefly, the bacterial colony was collected with the tip of a toothpick and directly used for PCR amplification. Quick Taq HS Dye mix was used as a PCR reagent (Toyobo, Tokyo, Japan). PCR primer sets, constructed for the predicted terminal ends of the plasmids, are listed in Supplementary Table 1. The thermal cycling conditions were set to: 94 °C for 3 min, 40 cycles of 94 °C for 30 s, 60 °C for 30 s, and 68 °C for 1 min 30 s.

### Growth kinetics and plasmid stability analysis

Bacterial strains were grown overnight in 5 mL THB. After diluting the bacterial culture to an absorbance (OD_620_) of 0.2, 1 μL of the dilution was added to 299 μL of THB and incubated at 37 °C for 24 h. The absorbance (OD_595_) was measured kinetically every 10 min in a Sunrise plate reader (Tecan, Männedorf, Switzerland) while incubating at 25 °C, 37 °C, or 42 °C without shaking. Growth rates (h) and fitness costs were determined using the Growth rate program (ver. 5.1) and the formula of Tedim *et al*.^50, 51^. Briefly, the growth rate (GR) of the pELF1-like plasmid-carrying strain was normalized by the GR of the pELF1-like plasmid non-carrying strain (relative to 1 [relative GR, RGR]). Plasmid fitness cost (%) was calculated as (1 – RGR) × 100. For both the growth curve and fitness cost analyses, we used nine biological replicates for each strain. Statistical analyses were performed using an unpaired *t*-test for growth rate and Mann–Whitney test for fitness cost.

The stability of the pELF1-like plasmid was evaluated through a 15-d passaging experiment in a liquid medium without selection pressure. Every 12 h, 5 µL of the bacterial culture was transferred to 5 mL of fresh medium and incubated continuously at 37 °C. After 5, 10, and 15 d, the number of pELF1-like plasmid-harboring and non-harboring strains was measured using vancomycin resistance as an indicator, and the ratio of pELF1-like plasmid-harboring cells to total cells was calculated. We conducted three independent biological experiments.

A carbon source-limited M1 medium was prepared as reported by Zhang *et al.*^52^. The bacteria were cultured in a THB medium and diluted to absorbance (OD_620_) 0.2. After centrifugation (13 000 × *g*, 3 min), the supernatant was discarded and the bacterial pellet was washed twice with PBS. The bacteria were inoculated with M1, M1 with 10 mM glucose, or M1 with 10 mM mannose and incubated at 37 °C for 24 h. The absorbance (OD_595_) was measured kinetically using the Sunrise plate reader (Tecan).

### Pulsed-field gel electrophoresis

PFGE was performed as previously described^7^. Briefly, a 1% agarose gel block embedded with enterococci was treated with lysozyme solution (10 mg/mL; Roche, Basel, Switzerland) at 37 °C for 6 h, followed by treatment with proteinase K solution (60 mAnson-U/mL; Merck Millipore, Darmstadt, Germany) at 50 °C for 48 h. After washing the treated plugs, they were subjected to PFGE using a CHEF Mapper (Bio-Rad Laboratories, Richmond, CA, USA) according to the manufacturer’s instructions (pulse from 5.3 to 66 s during 19.5 h at 6.0 V/cm and 4 °C).

### Plasmid copy number

The copy number of pELF1-like plasmids was calculated using qPCR as described by San Milan *et al*.^53^. Briefly, the copy number per chromosome was calculated as (1 + E_c_)^Ctc^ / (1 + E_p_)^Ctp^ × S_c_ / S_p_, where E_c_ and E_p_ are the efficiencies of the chromosome and plasmid qPCR amplification (relative to 1), respectively; C_tc_ and C_tp_ are the threshold cycles of the chromosome and plasmid, respectively; and S_c_ and S_p_ are the amplicon size (bp) of the chromosome and plasmid, respectively. DNA was extracted from cells cultured in THB medium containing 5 mg/L vancomycin and used for analysis immediately after entering the stationary phase. Considering its proximity to *oriC* for each species, *gyrB* was used as internal control and the replication-protein gene as the target gene for the pELF1-like plasmid. We conducted four biological replicates of this experiment and used an unpaired *t*-test for statistical analysis. The primers used are listed in Supplementary Table 1.

### Whole-genome sequencing analysis of pELF1-like plasmid-carrying isolates

Total DNA were prepared using a Gentra Puregene Yeast/Bact. Kit according to the manufacturer’s protocol (Qiagen, Hilden, Germany). Short reads were obtained using a MiniSeq system (Illumina) with a High Output Reagent Kit (300 cycles). The library for sequencing (150-bp paired-end, insert size, 500–900 bp) was prepared using a Nextera XT DNA Library Prep Kit (Illumina). The DNA library for nanopore MinION was prepared using a SQK-RBK004 Rapid Barcoding Kit according to the manufacturer’s protocol and then sequenced on a Flow Cell (R9.4.1, Oxford Nanopore Technologies, Oxford, UK). The long reads were basecalled using Guppy (ver. 3.30) and were assembled *de novo* using Canu (ver. 2.1.1) and then polished by short reads using Pilon (ver. 1.20.1)^54^. To obtain functional annotations, the assembled sequences were submitted to the RAST and DFAST annotation pipelines^55, 56^.

For the core genome analysis, the genomes of all strains were annotated in Prokka (ver. 1.14.6)^57^ for homogeneity. The core genes were identified and aligned in Roary (ver. 3.13.0)^58^. From the results of the core genome analysis, the phylogenetic tree was constructed using RAxML (ver. 8.2.4) under the GTRGAMMA model and then visualized in the Figtree software (ver. 1.4.4, http://tree.bio.ed.ac.uk/software/figtree/)^59^. We used Phandango and genome matcher (ver. 3.0.2) for visualization of the bacterial data set and genome comparison^60, 61^. MLST, antibiotic-resistance genes, replicon-type, toxin–antitoxin system, virulence genes, and MGEs in whole-genome data were detected using Staramr (ver. 0.5.1, https://github.com/phac-nml/staramr), Resfinder (ver. 4.1 server), ABRicate (https://github.com/tseemann/abricate), Plasmidfinder (ver. 2.1 server), TASer (https://shiny.bioinformatics.unibe.ch/apps/taser/), Virulence finder (ver. 2.0), ISfinder, and MobileElementFinder (ver. 1.0.3)^10, 11, 14, 34, 45, 62–64^. For the pELF comparison, pairwise nucleotide identity was calculated using MAFFT (ver. 7.450). CAI was calculated using the “cai” function of the EMBOSS package^65^. The genes encoding the ribosomal proteins of each of the four *Enterococcus* spp. were used as a reference set^66^. SNP and mapping analyses were performed using QIAGEN CLC Genomic Workbench (v11.0.1) (Qiagen).

### Transcriptome analysis

Bacteria used for RT-qPCR and RNA-seq were incubated overnight in BHI medium at 37 °C. The cultures were diluted 1,000-fold in fresh BHI medium and incubated until the late exponential phase. After treatment with RNA Protect Bacterial Reagent (Qiagen), RNA extraction was performed from 1 mL of the bacterial solution using an RNeasy Mini kit (Qiagen) according to the manufacturer’s protocol. For RT-qPCR analysis, cDNA was synthesized using the PrimeScript RT Master Mix (Takara Bio, Shiga, Japan), and real-time PCR was performed using the Luna Universal qPCR Master Mix (New England Biolabs, Ipswich, MA, USA) on an ABI 7500 Fast RT-PCR system (Thermo Fisher, Waltham, MA, USA). The primers used in this analysis are listed in Supplementary Table 1. For the RNA-seq analysis, RNA samples were outsourced to Novogene (Novogene, Beijing, China) for library preparation and sequencing (https://jp.novogene.com/). After preparing the strand-specific libraries, they were sequenced using an Illumina NovaSeq 6000 sequencer.

RNA data analysis was conducted using the Galaxy platform^67^. Reads were mapped to BM4105RF/pELF2 nucleotide sequences (accession numbers: CP030110 and AP022343) using Bowtie2 (ver. 2.4.2)^68^, and the numbers of aligned reads were counted with FeatureCounts (ver. 2.0.1)^69^. DESeq2 (ver. 2.11.40.6) was used to determine the differentially expressed genes (DEGs)^29^. In this analysis, significance was set at an adjusted *p*-value < 0.05, and a log_2_ fold change of >1 and <–1 was considered upregulated and downregulated, respectively. For the GO analysis, we used GOseq (ver. 1.44.0)^31^. Comparative analysis of plasmid and chromosomal gene transcription levels was performed using TPM values based on Kallisto Quant RNA-seq data, corrected for plasmid copy number^70^. The statistical analysis of gene transcript abundances between chromosomes and pELF1-like plasmids was performed using the Kolmogorov–Smirnov test.

## Acknowledgment

We are grateful to Reika Kawabata-Iwakawa (Division of Integrated Oncology Research, Gunma University Initiative for Advanced Research (GIAR), Gunma University), Yohei Morishita, and Saori Fujimoto (Laboratory for Analytical Instruments, Education and Research Support Center, Gunma University Graduate School of Medicine) for their helpful discussions and assistance with RNA integrity measurement.

This study was supported by grants from the Japanese Ministry of Health, Labor and Welfare [Research Program on ensuring Food Safety: 21KA1004], grants from the Japan Agency for Medical Research and Development (AMED) [JP22fk0108604 and JP22wm0225008 to H. Tomita; JP22gm1610003, JP22fk0108133, JP22fk0108139, JP22fk0108642, JP22wm0225004, JP22wm0225008, JP22wm0225022, JP22wm0225022, JP22wm0325003, JP22wm0325037 to M. Suzuki], grants from the Ministry of Education, Culture, Sports, Science and Technology (MEXT), Japan [20K07509 and 21H03622 to M. Suzuki; Grant-in-Aid for Early-Career Scientists 22K16368 to Y. Hashimoto], and the grant from Gunma Foundation and Health Science [Research Grants Provided by Gunma Foundation and Health Science].

## Author contributions

Y.H.1, M.S., and H.T. conceived and designed the study. Y.H.1, S.K, Y.H.2, T.N., J.K., H.H., and K.T. collected bacterial isolates and performed bacterial characterizations. Y.H.1 and M.S. performed the collection and analysis of WGS and transcriptome data. Y.H.1 conducted all other efforts. H.T. supervised the study. All authors read and approved the manuscript for publication. (Y.H.1; Yusuke Hashimoto, Y.H.2; Yuki Hirahara)

