## Supplemental Figures for "Enterococcal linear plasmids adapt to *Enterococcus faecium* and spread within multidrug-resistant clades"

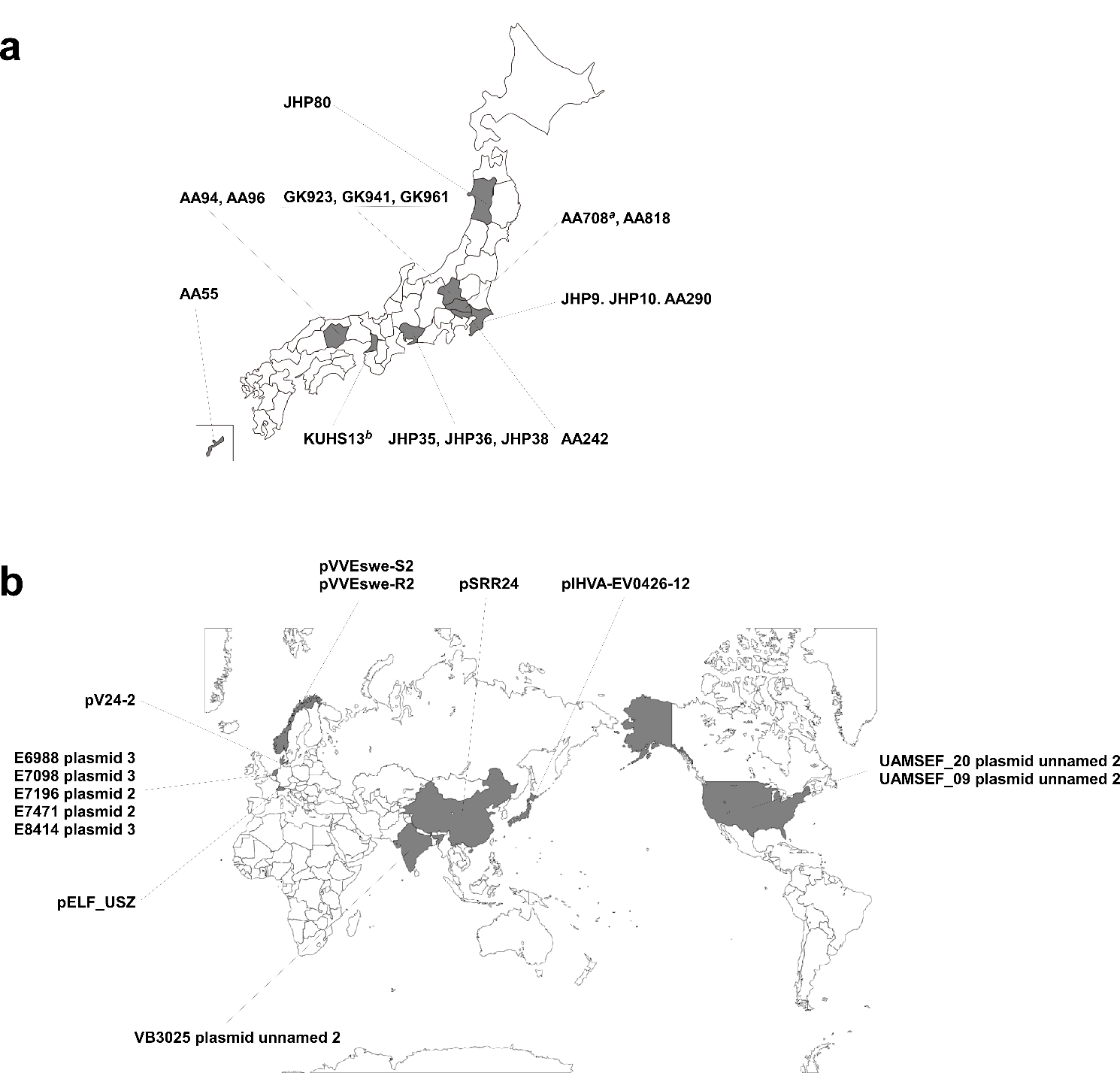

**Supplementary Figure 1. Geographic areas of detection of enterococci harboring pELF1-like plasmids.**

(a) Prefectures where enterococci harboring pELF1-like plasmids were isolated in Japan, are shown in black. *^a,b^* AA708 and KUHS13 strain was a strain previously reported. Underlined strains indicate VSE strains. (b) The Countries where presumed pELF1-type plasmid harboring strains are detected are shown in black.

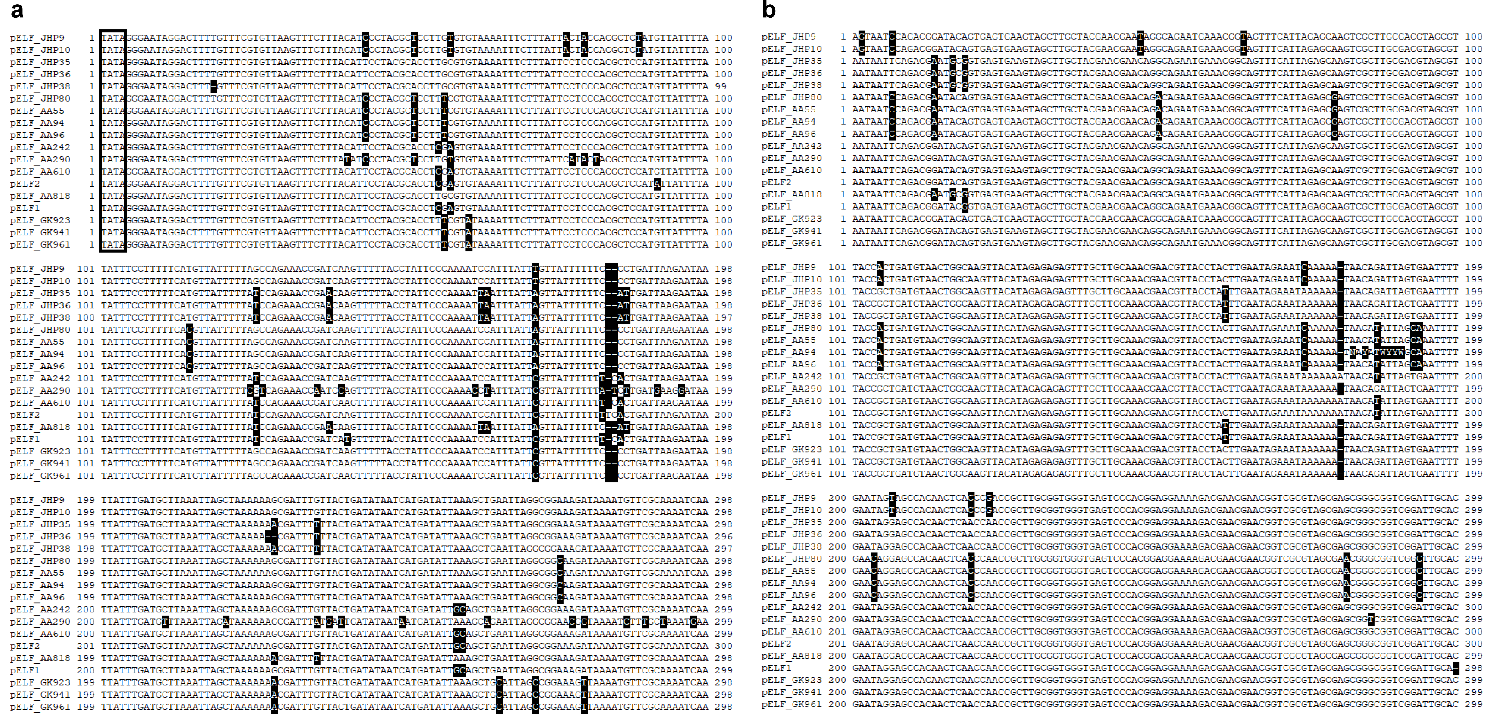

**Supplementary Figure 2. Terminal nucleotide sequence comparisons of pELF-like plasmids.**

The nucleotide sequences at the ends of the pELF-like plasmids were aligned and compared using ClustalW. (a) Sequence comparison of the hairpin end, (b) sequence comparison of the invertron end. Sequence mismatches are indicated by the black background. The -TATA- sequence forming the hairpin end is circled by a square (a).

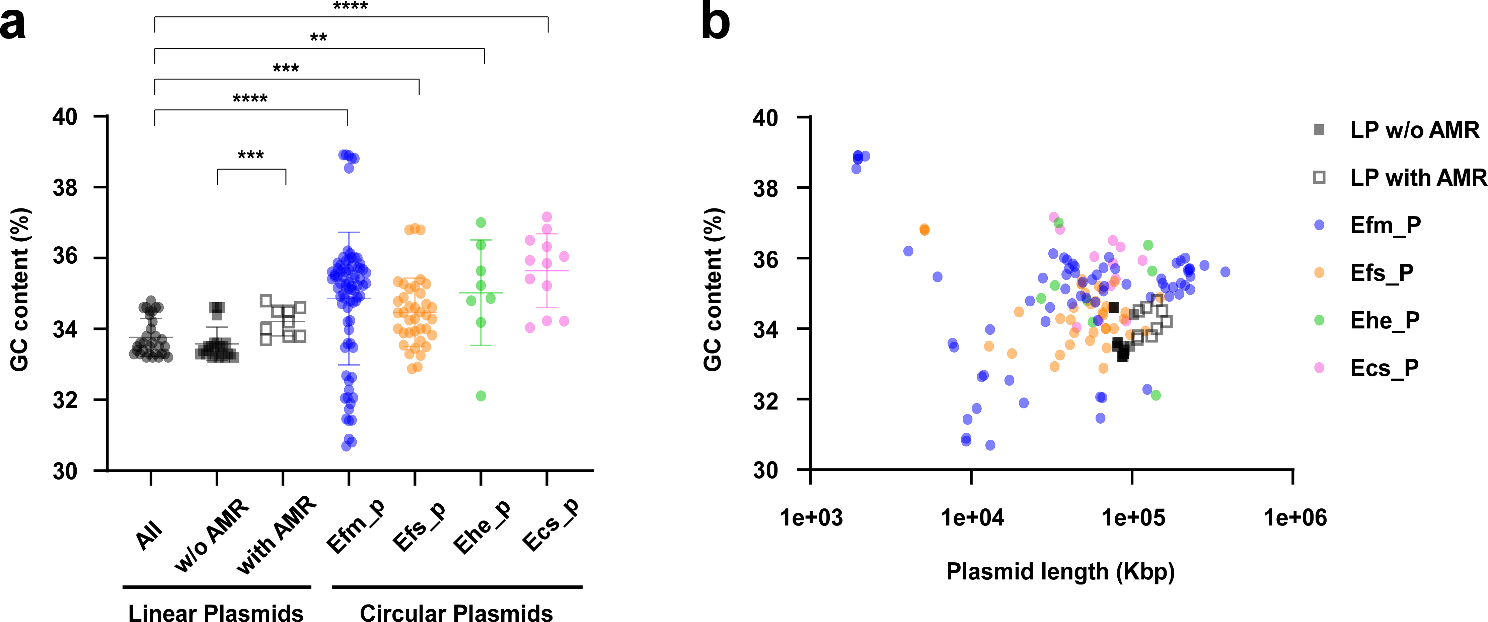

**Supplementary Figure 3. Scatter plot of GC content and plasmid size of enterococcal circular plasmids along with 32 pELF1-like plasmids.**

The GC content of 138 circular plasmids and 32 pELF1-like plasmids was analyzed. (a) The GC content of the plasmids possessed by each species and (b) the size and GC content of the plasmids are shown in the scatter plots. Black dots indicate All linear plasmids, Black squares indicate pELF1-like plasmids without AMR regions, white squares indicate linear plasmids with AMR regions, blue dots indicate circular plasmids of *E. faecium*, orange dots indicate circular plasmids of *E. faecalis*, green dots indicate circular plasmids of *E. hirae*, and pink dots indicate circular plasmids of *E. casseliflavus*. All circular plasmids were checked for possession of *rep* genes using PlasmidFinder (2.1). ****, *p* <0.0001; ***, *p* <0.001; **, *p* <0.01 (Mann–Whitney test). LP stands for pELF1-like linear plasmids, Efm_p for circular plasmids possessed by *E. faecium*, Efs_p for circular plasmids possessed by *E. faecalis*, Ehe_p for circular plasmids possessed by *E. hirae*, and Ecs_p for circular plasmids possessed by *E. casseliflavus*.

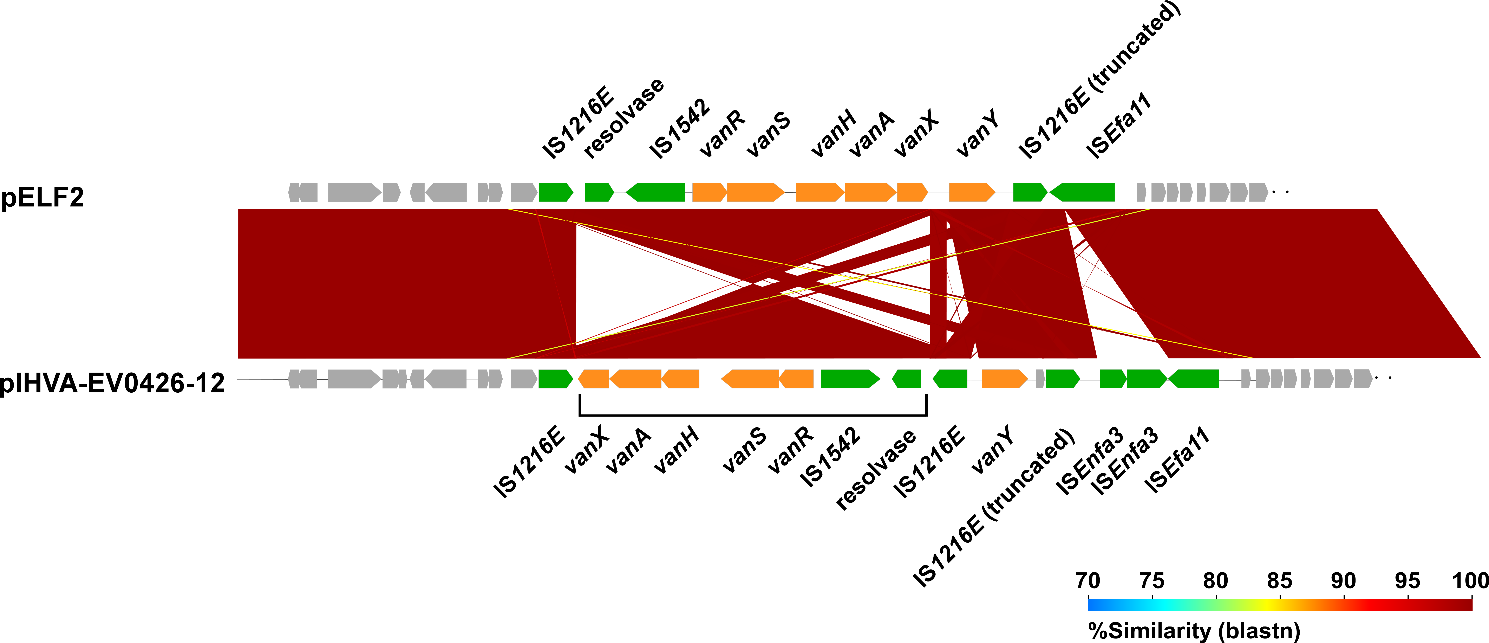

**Supplementary Figure 4. Comparative analysis of the structures around *vanA* gene cluster of pELF2 and pIHVA-EV0426-12.**

The inverted region of pIHVA-EV0426-12 compared to pELF2 was shown. The panel shows the genetic structure. The green panels represent mobile genetic element-related genes, and the orange panels represent AMR genes.

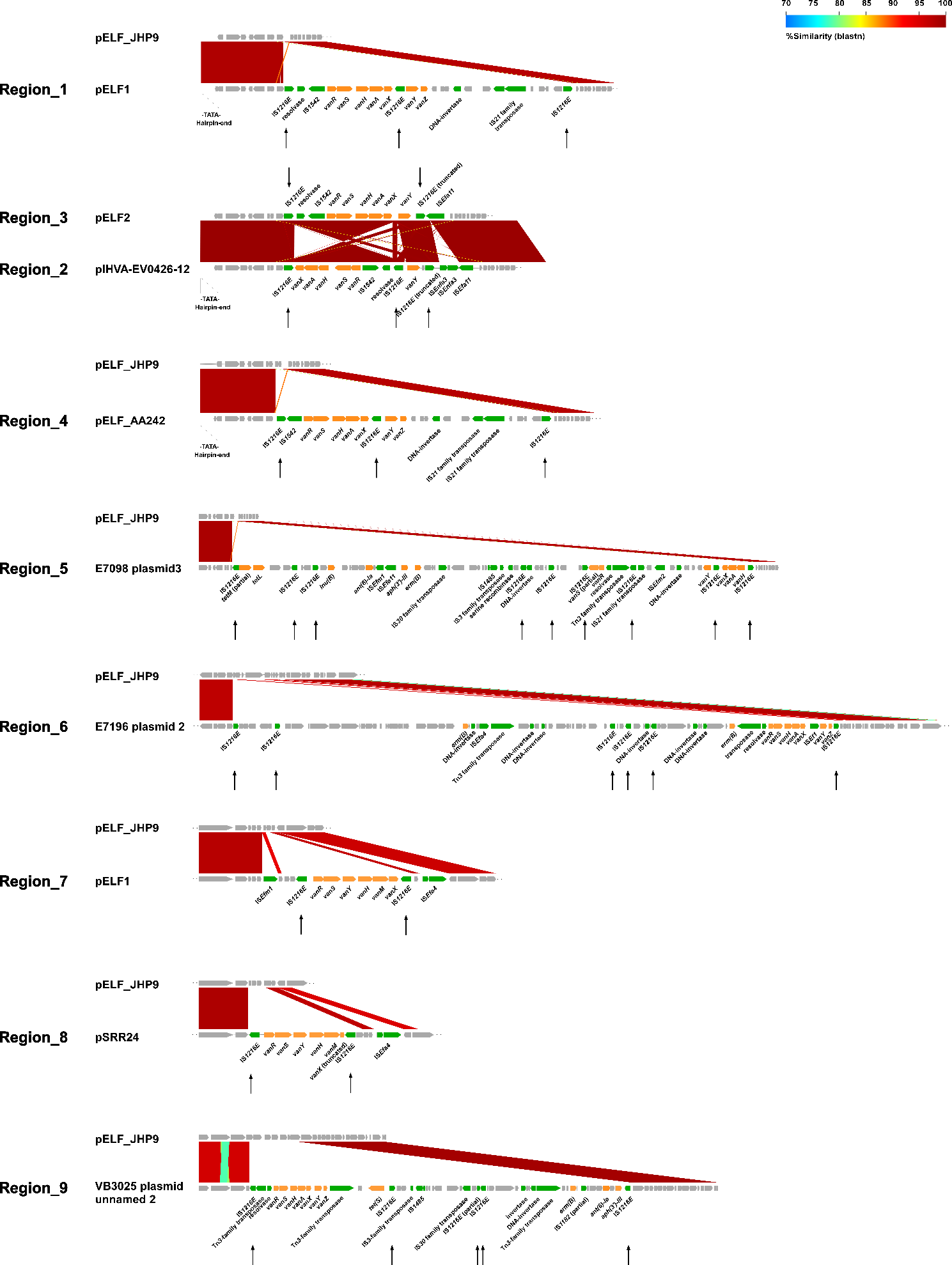

**Supplementary Figure 5. Comparison of AMR regions containing vancomycin resistance genes in pELF1-like plasmids.**

The nine AMR regions containing vancomycin resistance genes on the pELF1-like plasmid were compared. The numbers assigned to each AMR region are consistent with those in Figure 2. The panels show the genetic structures. The green panels represent mobile genetic element-related genes, and the orange panels represent AMR genes. The arrows pointing vertically to the panels indicate the position of IS*1216E*.

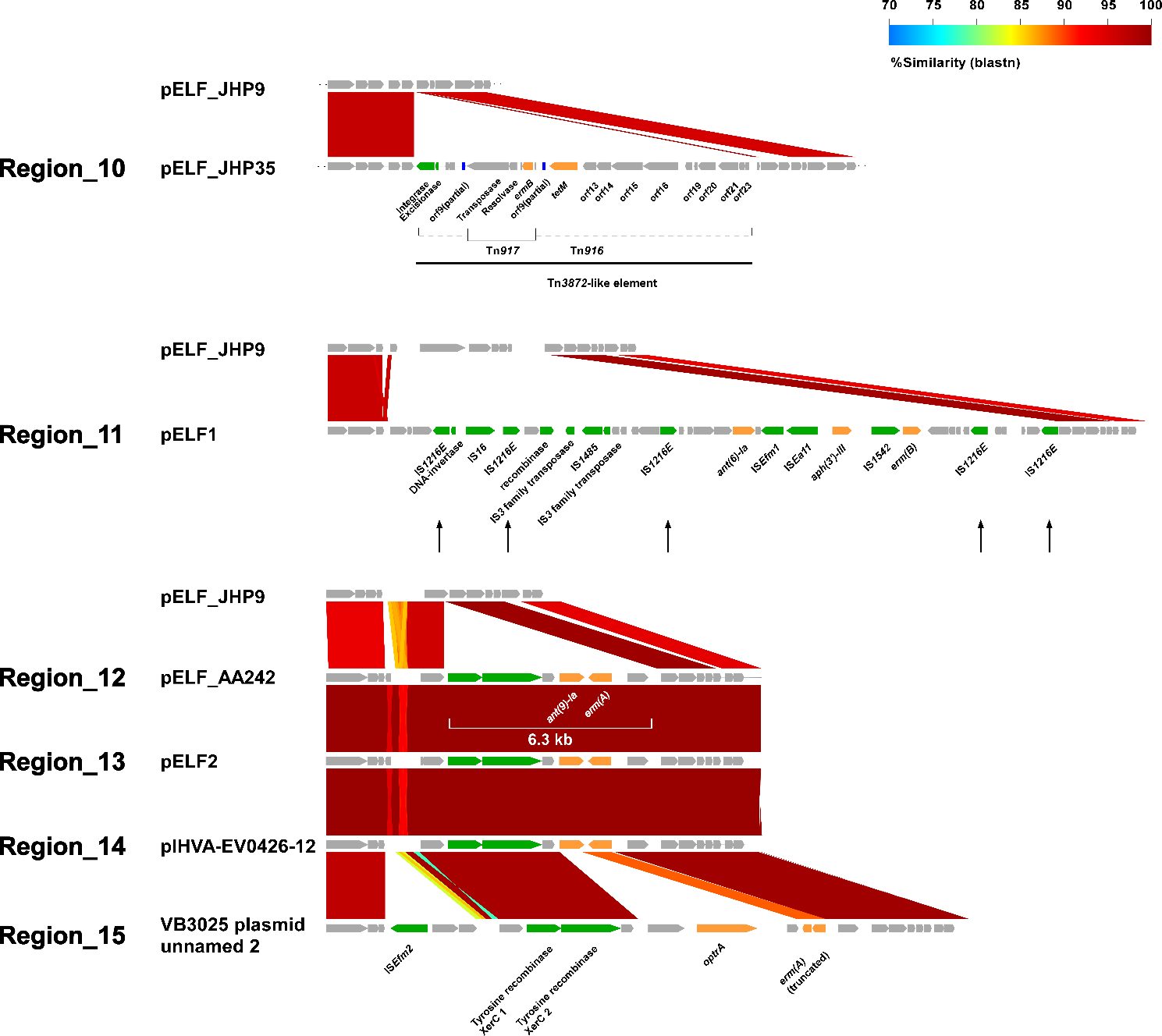

**Supplementary Figure 6. Comparison of AMR regions without vancomycin resistance genes in pELF1-like plasmids.**

The nine AMR regions without vancomycin resistance genes on the pELF1-like plasmid were compared. The numbers assigned to each AMR region are consistent with those in Figure 2. The panel shows the genetic structure. The green panels represent mobile genetic element-related genes, and the orange panels represent AMR genes. The arrows pointing vertically to the panels indicate the position of IS*1216E*.

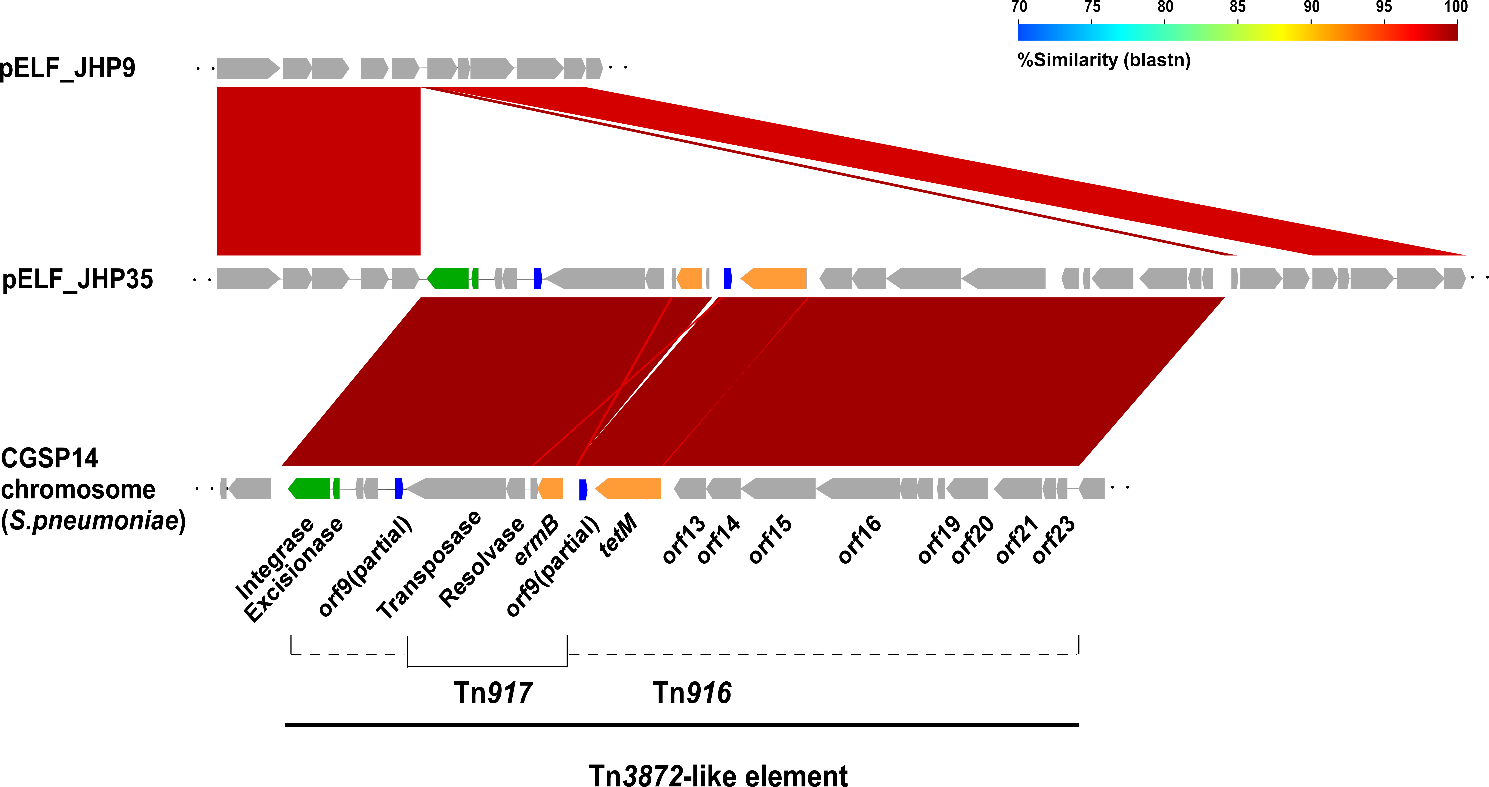

**Supplementary Figure 7. Comparative analysis of Tn*3872*-like element in pELF_JHP35.**

The structures of the *Tn3872*-like element of pELF_JHP35 and (B) the regions around *ant(9)-Ia* and *erm*(A) of pELF_AA242 were compared with those of pELF_JHP9. The blue panels represent partial *orf9* disrupted by Tn*917*, the orange panels represent AMR genes, and the green panels represent mobile genetic element-related genes.

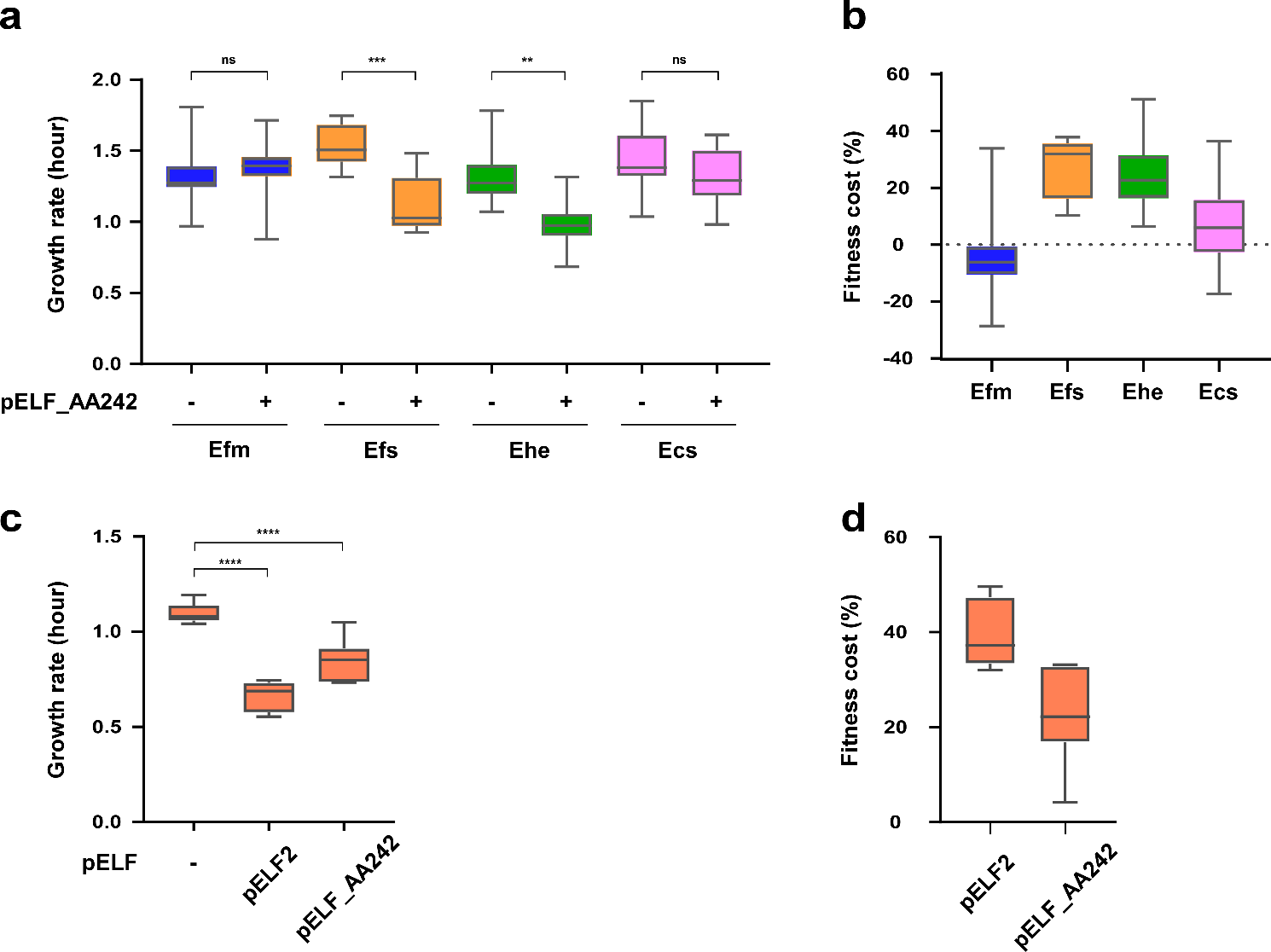

**Supplementary Figure 8. Additional analysis of the effect of pELF1-like plasmids on the growth of enterococci.**

The effects of possession of pELF1-like plasmid (pELF_AA242) on (a) the growth rate and (b) the fitness cost of four Enterococcus species are shown. Efm: *E. faecium* (BM4105RF), Efs: *E. faecalis* (FA2-2), Ehe: *E. hirae* (ATCC9790RF), Ecs: *E. casseliflavus* (KT06RF). The box-and-whiskers indicate the minimum to maximum values. The unpaired *t*-test was used to analyze the pELF1-like plasmid harboring and non- harboring strains in the growth rate ((a) *p* = 0.6564, *p* = 0.0002, *p* = 0.0016, and *p* = 0.2744, respectively, and (c) *p* <0.0001, and *p* <0.0001). The *E. faecalis* strain was changed to OG1RF, and (c) the growth rates and (d) the fitness costs of pELF2 and pELF242 were analyzed. For this analysis of growth rate and fitness cost, at least six biological replicates were subject to the analyses in duplicate. **, *p* ≤0.01; ***, *p* ≤0.001; ****, *p* ≤0.0001; ns, not significant.

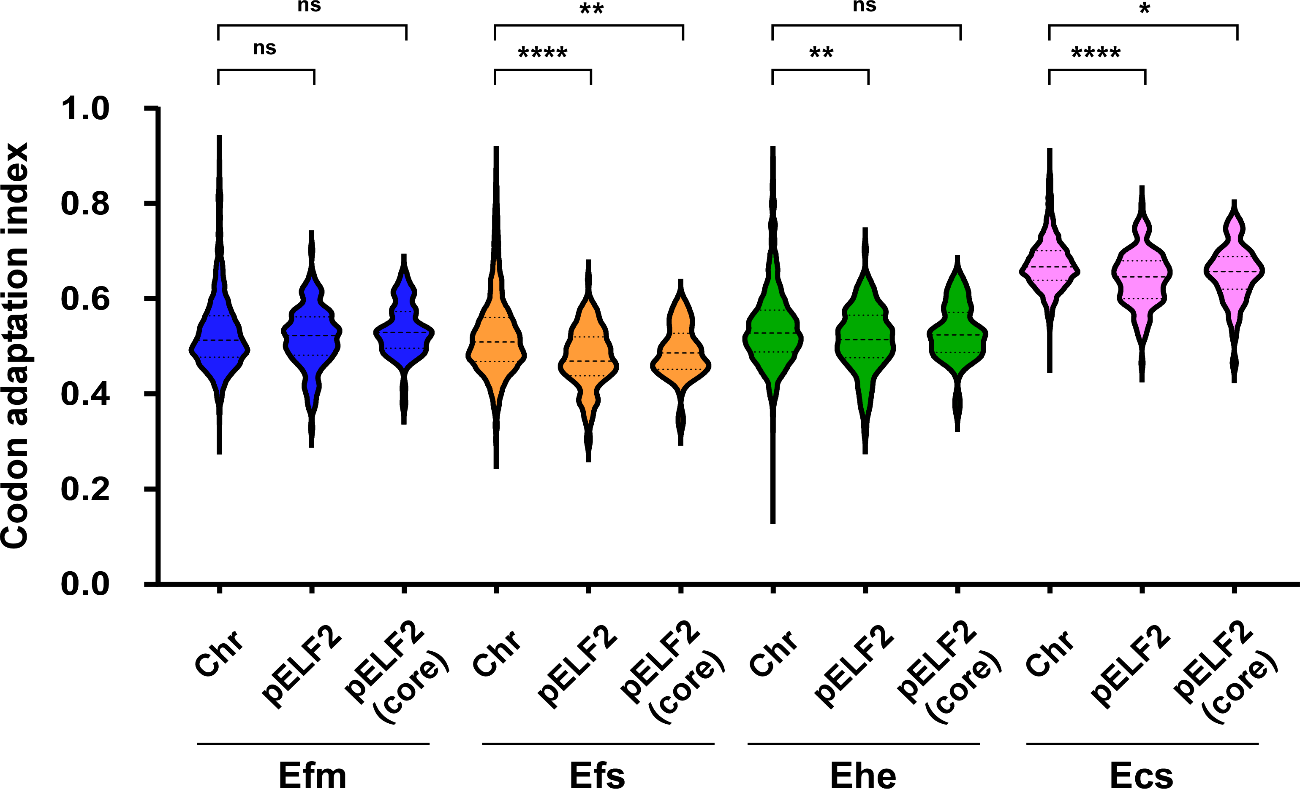

**Supplementary Figure 9. Codon adaptation index analysis.**

For the four enterococci species, the codon adaptation index for the chromosomal genes (Chr), the genes on pELF2, and the core genes on pELF2 were determined and illustrated in violin plots. *, *p* ≤ 0.05; **, *p* ≤ 0.01; ****, *p* ≤ 0.0001 (Mann–Whitney test). Efm: *E. faecium* (BM4105RF), Efs: *E. faecalis* (FA2-2), Ehe: *E. hirae* (ATCC9790RF), Ecs: *E. casseliflavus* (KT06RF).

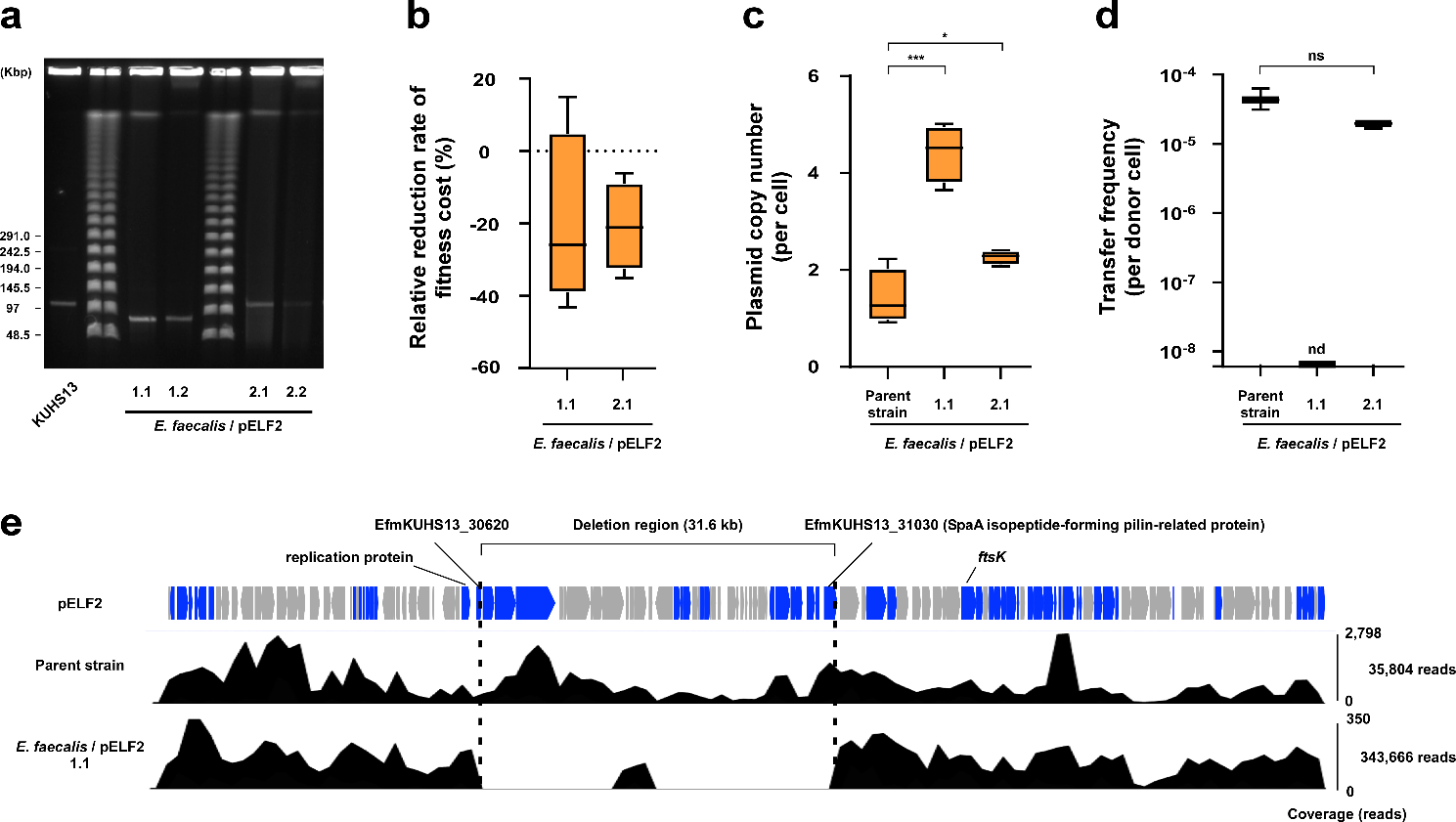

**Supplementary Figure 10. Analysis of structure and stability of the pELF1-like plasmid in *E. faecalis.***

(a) PFGE analysis was conducted to assess the size of the pELF1-like plasmid (pELF2) harbored by *E. faecalis* (FA2-2) on day 5 of the long-term passage assay (day 5-evolved strains) (Figure 3C). Colonies of day 5-evolved strains were randomly selected and subjected to PFGE (S1 nuclease untreated) along with the wild strain harboring pELF2 (KUHS13; accession number: SAMD00202474). (b) The percentage reduction in fitness cost relative to the parent strain (FA2-2/pELF2) was calculated in quadruplicate. (c) Plasmid copy number was measured using qPCR. Quadruplicate results were shown in the box-and-whisker diagram. *; *p* ≤0.05, ***; *p* ≤ 0.001 (unpaired *t*-test; *p* = 0.0004, and *p* = 0.0286). (d) Day 5-evolved strains were subjected to filter mating with BM4105SS as the recipient strain. The plasmid transfer frequencies were calculated and shown in a box-and-whisker diagram. ns, not significant; nd, not detected (Mann–Whitney test). (e) A short read set of the Day 5-evolved strain (1.1), as well as the parent strain, were mapped to the pELF2 nucleotide sequence, and the 31.6 kb deletion region in the Day 5-evolved strain (1.1) was shown.

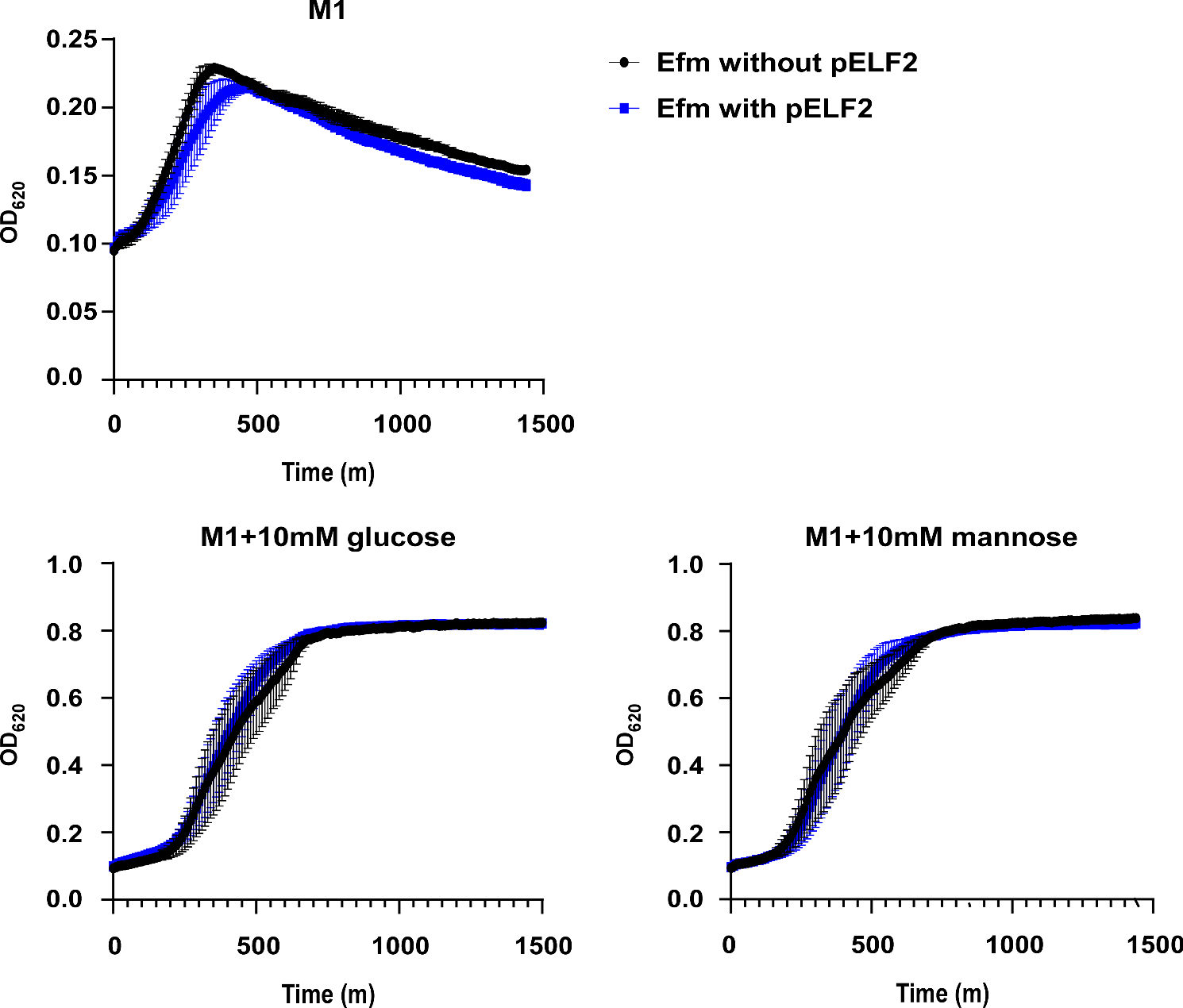

**Supplementary Figure 11. Effect of pELF1-like plasmid carriage on *E. faecium* growth in M1 medium.**

The impact of pELF1-like plasmid (pELF2) on the growth of *E. faecium* (BM4105RF) under carbon source-limited culture conditions was analyzed. Black dots and lines indicate strains without pELF2, and blue dots and lines indicate strains with pELF2. The mean and standard error are plotted in four independent experiments. Individual culture conditions are shown at the top of each graph.

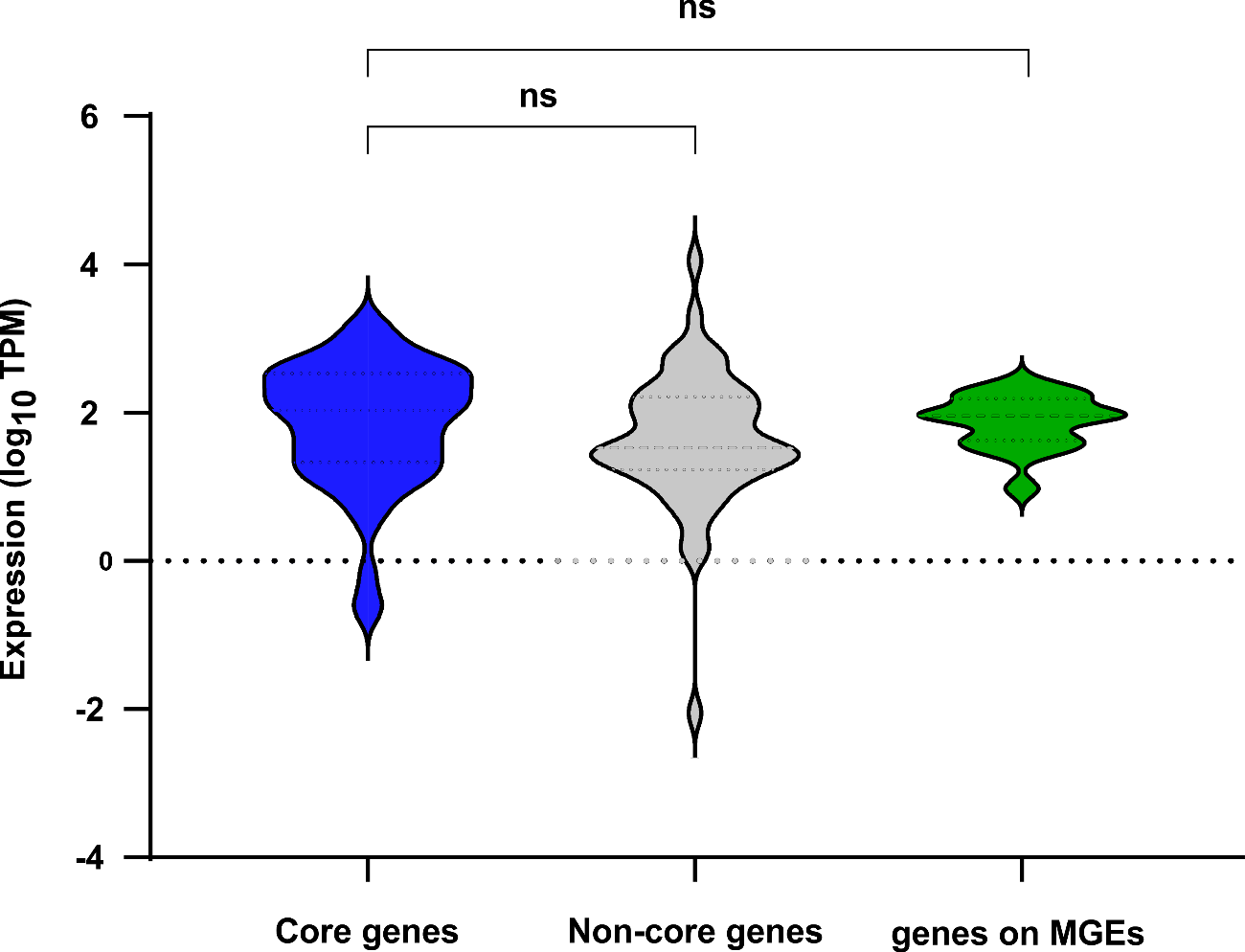

**Supplementary Figure 12. Comparative analysis of transcript levels of genes on the pELF1-like plasmid.**

Transcription levels (TPM) of genes on pELF2 were compared by classification of core genes, non-core genes, and accessory genes (MGE) and illustrated in violin plots. ns, not significant (Mann–Whitney test).
